# During *Aspergillus* infection, neutrophil, monocyte-derived DC, and plasmacytoid DC enhance innate immune defense through CXCR3-dependent crosstalk

**DOI:** 10.1101/2020.05.05.079517

**Authors:** Yahui Guo, Shinji Kasahara, Anupam Jhingran, Nicholas L. Tosini, Bing Zhai, Mariano A. Aufiero, Kathleen A.M. Mills, Mergim Gjonbalaj, Vanessa Espinosa, Amariliz Rivera, Andrew D. Luster, Tobias M. Hohl

**Affiliations:** Infectious Disease Service, Department of Medicine, Memorial Sloan Kettering Cancer Center, New York, NY, USA; Immunology Program, Memorial Sloan Kettering Cancer Center, New York, NY, USA; Louis V. Gerstner Jr. Graduate School of Biomedical Sciences, Sloan Kettering Institute, Memorial Sloan Kettering Cancer Center, New York, NY, USA; Immunology and Microbial Pathogenesis Graduate Program, Weill Cornell Graduate School, New York, NY, USA; Center for Immunity and Inflammation, Rutgers Biomedical and Health Sciences (RBHS), Newark, NJ, USA; Department of Pediatrics, New Jersey Medical School, Rutgers Biomedical and Health Sciences (RBHS), Newark, NJ, USA; Center for Immunology and Inflammatory Diseases, and Division of Rheumatology, Allergy, and Immunology, Massachusetts General Hospital, Charlestown, MA, USA

**Keywords:** *Aspergillus fumigatus*, CXCR3, CXCL9, CXCL10, plasmacytoid DC, monocyte, neutrophil, dendritic cell, innate immunity, cytokine, lung, fungus, crosstalk

## Abstract

*Aspergillus fumigatus*, a ubiquitous mold, is a common cause of invasive aspergillosis (IA) in immunocompromised patients. Host defense against IA relies on lung-infiltrating neutrophils and monocyte-derived dendritic cells (Mo-DCs). Here, we demonstrate that plasmacytoid dendritic cells (pDCs), which are prototypically anti-viral cells, participate in innate immune crosstalk underlying mucosal antifungal immunity. *Aspergillus*-infected murine Mo-DCs and neutrophils recruited pDCs to the lung by releasing the CXCR3 ligands, CXCL9 and CXCL10, in a Dectin-1/Card9- and type I and III interferon-signaling dependent manner, respectively. During aspergillosis, circulating pDCs entered the lung in response to CXCR3-dependent signals. Via targeted pDC ablation, we found that pDCs were essential for host defense in the presence of normal neutrophil and Mo-DC numbers. Although interactions between pDC and fungal cells were not detected, pDCs regulated neutrophil NADPH oxidase activity and conidial killing. Thus, pDCs act as positive feedback amplifiers of neutrophil effector activity against inhaled mold conidia.

## Introduction

*Aspergillus fumigatus* forms airborne spores (conidia) that humans typically clear in a silent and asymptomatic manner. Due to the growing number of humans that live in immune compromised states, *A. fumigatus* is the most common and lethal agent of mold pneumonia worldwide (Brown et al., 2012; Latge and Chamilos, 2019; Lionakis and Levitz, 2018; Tischler and Hohl, 2019). In humans and mice, sterilizing immunity against *A. fumigatus* conidia depends on the action of myeloid cells, primarily neutrophils, lung-infiltrating monocytes and monocyte-derived DCs (Mo-DCs), as well as alveolar macrophages; all lymphoid cells are redundant for host defense (Espinosa et al., 2014; Hohl et al., 2009; Latge and Chamilos, 2019; Mircescu et al., 2009). Phagocyte NADPH oxidase is central to fungal clearance, as evidenced by a 40-55% lifetime prevalence of IA in patients with chronic granulomatous disease (Marciano et al., 2015). Reactive oxygen species induce a regulated cell death process in neutrophil-engulfed conidia, and thereby protect the respiratory tract from tissue-invasive hyphae and fungal dissemination (Shlezinger et al., 2017).

Using targeted cell depletion strategies based on murine CCR2 promoter-dependent diphtheria toxin receptor (DTR) transgene expression (Espinosa et al., 2014; Hohl et al., 2009), recent studies implicated CCR2^+^ inflammatory monocytes and Mo-DCs in sterilizing antifungal immunity, both by direct fungal killing and by regulating the lung inflammatory milieu (Espinosa et al., 2017; Espinosa et al., 2014). Since depletion of CCR2^+^ cells diminished neutrophil antifungal activity, two models could explain these findings. First, CCR2^+^ monocytes and Mo-DCs could release mediators that act directly on neutrophils to enhance antifungal effector functions. Second, CCR2^+^ monocytes and Mo-DCs could mediate the recruitment or activation of a third cellular constituent that conditions the lung inflammatory milieu independent of contributions from CCR2^+^ monocytes and their derivatives. Here, we provide evidence for the second model and identify pDCs as a third leukocyte constituent that is essential for innate immune crosstalk and host defense against *A. fumigatus* in an otherwise immune competent host.

The mechanism by which pDCs contribute to antifungal immunity in the lung remains an open question. *In vitro*, human pDCs may spread over *A. fumigatus* hyphae to blunt fungal metabolic activity, and, in rare instances, undergo a cell death process that may result in extracellular traps (Loures et al., 2015; Ramirez-Ortiz et al., 2011). Mice treated with a mAb (i.e., *α*-PDCA-1/CD317) that primarily targets pDCs in the steady state, but likely targets additional leukocyte subsets under inflammatory conditions (Blasius et al., 2006), are susceptible to *A. fumigatus* challenge. In this study, we integrate pDCs into a model of innate immune crosstalk that is critical for defense against *A. fumigatus* in the lung. We found that fungus-infected Mo-DCs and neutrophils utilize Dectin-1/Card9 signaling to release CXCL9 and responded to type I and type III interferon signaling to release CXCL10. These CXCR3 ligands promoted pDC egress from the circulation into the infected lung. Lung CXCR3^+^ pDCs enhanced neutrophil NADPH oxidase activity and fungal killing, preventing the formation of tissue-invasive hyphae and promoting sterilizing immunity. These findings integrate antifungal pDCs into a model of mucosal immune defense against inhaled molds.

## Results

### Mo-DCs and neutrophils produce CXCL9 and CXCL10 during *A. fumigatus* infection

To examine specific contributions of CCR2^+^ monocytes and Mo-DCs to the lung inflammatory milieu during *A. fumigatus* challenge and to eliminate potential contributions of CCR2^+^ NK cell and CD8^+^ T cell subsets, we crossed the CCR2-diphtheria toxin receptor (DTR) transgene to Rag2^-/-^ interleukin-2 receptor *γ* chain^-/-^ mice. Lung homogenates of DT-treated CCR2-DTR^+/-^ Rag2^-/-^ interleukin-2 receptor *γ* chain^-/-^ mice (CCR2-DTR^+/-^ *Rag2*^*-/-*^*Il2rg*^*-/-*^) contained less CXCL10 (IP-10), CCL5 (RANTES), CCL17 (TARC), and CCL20 (MIP-3*α*) compared to DT-treated non-transgenic *Rag2*^*-/-*^*Il2rg*^*-/-*^ littermates at 36 h post-infection (pi), as measured in a cytokine array (Figures S1A and S1B).

Because a *CXCL10* polymorphism is implicated in IA susceptibility in hematopoietic cell transplant recipients (Fisher et al., 2017; Mezger et al., 2008), we measured lung levels of CXCL9 and CXCL10, both CXCR3 ligands, in naïve and in fungus-infected C57BL/6 mice. *A. fumigatus* challenge induced CXCL9 and CXCL10 expression in the lung, with a peak at 48 h pi (Figure 1A). To validate the multiplex array data, we ablated CCR2^+^ monocytes and Mo-DCs in CCR2-DTR^+/-^ *Rag2*^*-/-*^*Il2rg*^*-/-*^ mice and observed a 50-70% reduction in lung CXCL9 and CXCL10 levels at 48 h pi (Figure S1C). Ablation of CCR2^+^ monocytes and Mo-DCs in conventional CCR2-DTR^+/-^ (CCR2 Depleter) mice that contain lymphoid lineage cells yielded nearly identical results (Figures 1B and 1C), consistent with the model that CCR2^+^ myeloid cells represent a major cellular source of CXCR3 ligands during acute *A. fumigatus* infection.

**Figure 1.**
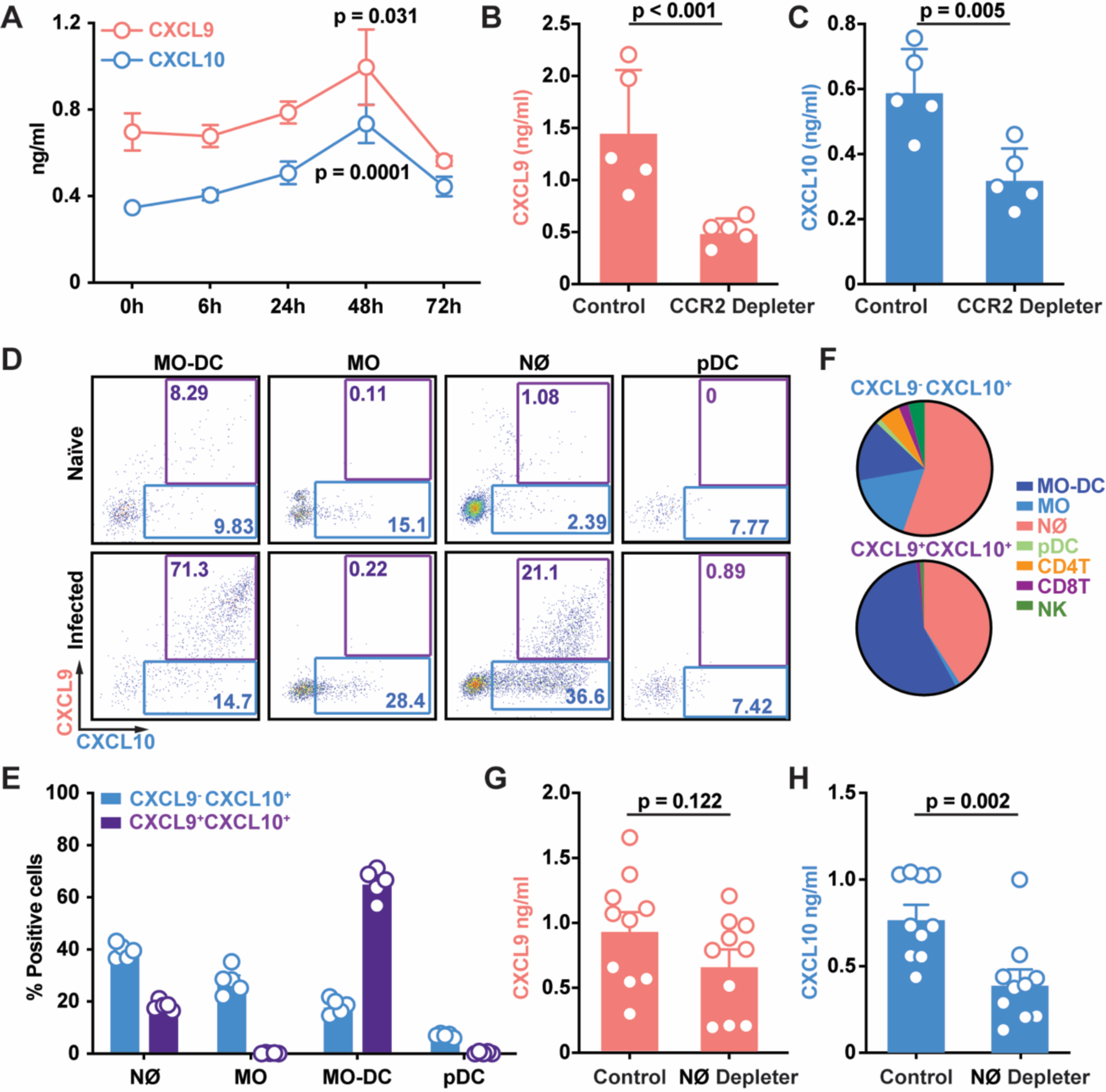
CCR2^+^ Mo-DCs and neutrophils produce CXCL9 and CXCL10 during *A. fumigatus* challenge. (A-C) Lung CXCL9 and CXCL10 levels in (A) C57BL/6 mice (n = 5) at 0-72 h pi or (B-C) in DT-treated CCR2-DTR^+/-^ and Non-Tg (CCR2-DTR^-/-^) littermates (n = 5) at 48 h pi with 3 × 10^7^ CEA10 conidia. (D) Representative plots of RFP (CXCL9) and BFP (CXCL10) expression in indicated lung leukocytes isolated from Rex3 Tg → C57BL/6.SJL BM chimeric mice at baseline (naïve, top row) and 48 h pi with 3 × 10 ^7^ CEA10 conidia (infected, bottom row). The blue and purple gates indicate the frequency of BFP^+^ (CXCL9^+^) and BFP^+^RFP^+^ (CXCL9^+^ CXCL10^+^) cells, respectively. (E) The graphs indicate the frequency of CXCL9^-^ CXCL10^+^ and CXCL9^+^ CXCL10^+^ neutrophils, monocytes, Mo-DCs, and pDCs and (F) the cellular identity of all CXCL9^-^ CXCL10^+^ (top) and CXCL9^+^ CXCL10^+^ lung leukocytes (bottom) at 48 h pi. (G-H) Lung CXCL9 and CXCL10 levels in DT-treated ROSA26-iDTR^Mrp8-Cre^ → C57BL/6.SJL (Neutrophil Depleter) or non-Cre iDTR littermates → C57BL/6.SJL (Control) BM chimeric mice (n = 10) at 48 h pi with 3-4 × 10^7^ heat-killed swollen CEA10 conidia. Data from 2 independent experiments were pooled. (A-C, G, H) Dots represent individual mice and data were presented as mean ± SEM. Statistical analysis: (A) Kruskal-Wallis test, timepoints compared to t = 0 h, (B, C, G, H) Mann-Whitney test. See also Figure S1.

To visualize hematopoietic cellular sources of CXCL9 and CXCL10, we generated chimeric mice (REX3 Tg → C57BL/6) in which radiosensitive hematopoietic cells encoded RFP and BFP transgenes that were driven by the *Cxcl9* and *Cxcl10* promoters, respectively (Groom et al., 2012). Following *A. fumigatus* challenge, Mo-DCs were the major cell type that expressed the RFP and BFP transgenes, consistent with the CCR2^+^ cell ablation data in the CCR2-DTR^+/-^ *Rag2*^*-/-*^*Il2rg*^*-/-*^ and CCR2-DTR^+/-^ backgrounds (Figures 1D, 1E and 1F). The majority of Mo-DCs produced CXCL9 and CXCL10 simultaneously, while a minority were positive only for CXCL10 (Figures 1E and 1F). Monocytes expressed primarily the CXCL10 transgene, while CXCL9 transgene expression was undetectable at baseline and during fungal infection. Neutrophils expressed both reporter transgenes during respiratory fungal infection, albeit with a lower frequency compared to Mo-DCs. pDCs, CD4^+^ and CD8^+^ T cells, and NK cells expressed very low levels of the reporter transgenes, respectively (Figures 1D-1F, S1D and 1E).

To ascertain that neutrophils contribute to CXCL9 and CXCL10 production, we generated BM chimera mice that enabled DT-induced neutrophil depletion (ROSA26-iDTR^Mrp8-Cre^ → C57BL/6.SJL mice, neutrophil Depleter) and (non-Cre iDTR littermates → C57BL/6.SJL), and measured CXCL9 and CXCL10 levels following challenge with heat-killed swollen *A. fumigatus* conidia 48 h pi. This experimental set-up was utilized to eliminated potential differences in fungal growth observed in neutrophil depletion studies (Bonnett et al., 2006; Espinosa et al., 2014; Mehrad et al., 1999; Mircescu et al., 2009). Neutrophil-depleted mice exhibited significantly lower CXCL10 lung levels compared to non-depleted littermates, though CXCL9 lung levels were similar (Figures 1G and 1H). Although CCR2^+^ Mo-DCs were the major cellular source of CXCL9 and CXCL10 during acute *A. fumigatus* challenge, neutrophils contributed to CXCL10 production as well.

### Distinct signaling pathways promote CXCL9 and CXCL10 production by fungus-infected myeloid cells in the lung

To examine whether fungal uptake by Mo-DCs and neutrophils drives CXCL9 and CXCL10 expression, we infected chimeric REX3 Tg → C57BL/6 mice with Alexa 633 (AF633)-labeled conidia and compared RFP and BFP expression in fungus-engaged (AF633^+^) and bystander (AF633^-^) leukocytes. Fungus-engaged Mo-DCs and neutrophils had much higher levels of fluorescent transgene expression than corresponding bystander cells, indicating that fungal uptake promotes Mo-DC CXCL9 and CXCL10 production, neutrophil CXCL10 production (Figures 2A-2E), and, to a lesser extent, monocyte CXCL9 production (Figures S2A, 2B).

**Figure 2.**
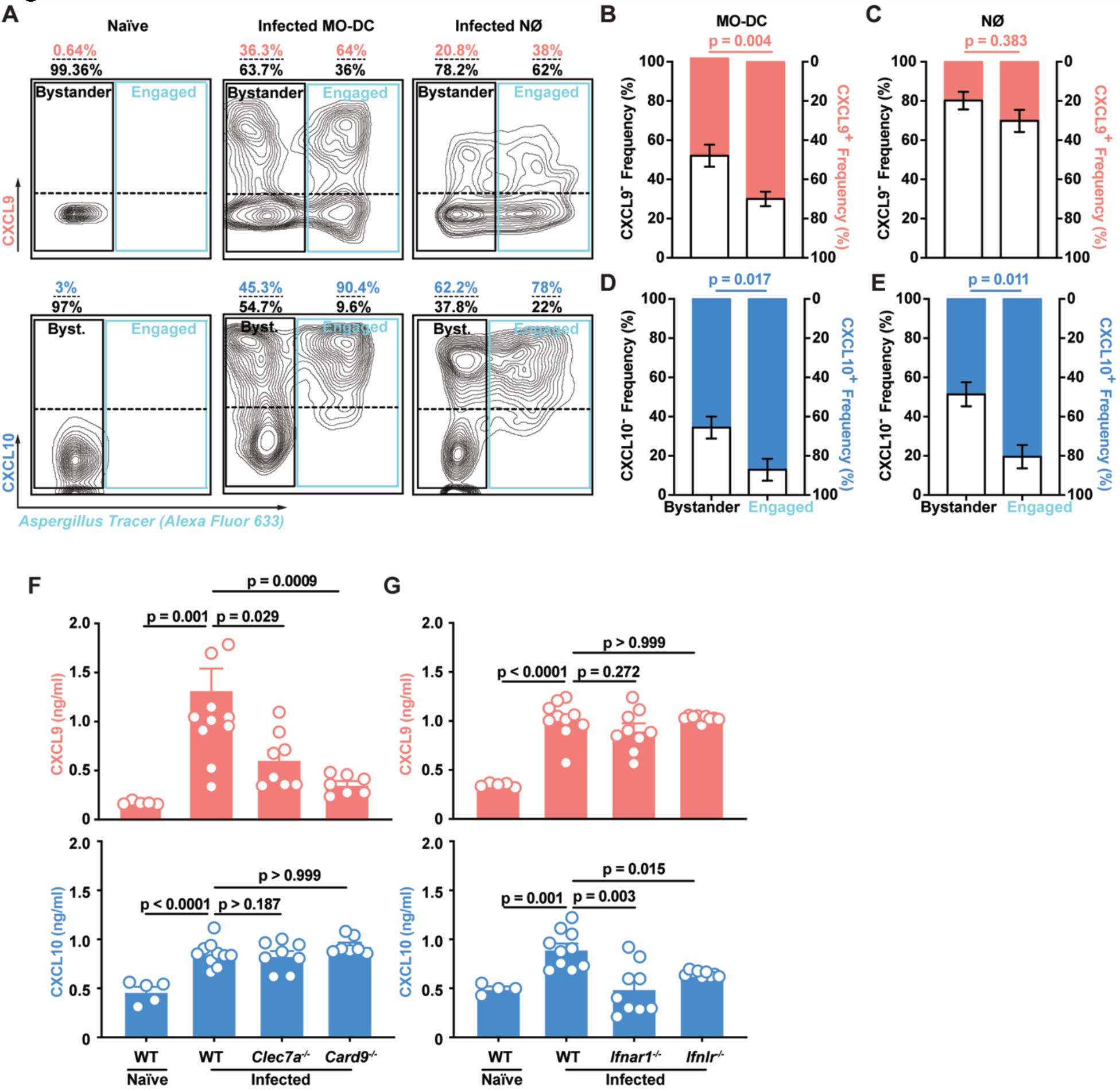
Fungal Uptake and Dectin-1/Card9 and IFN signaling regulate lung CXCL9 and CXCL10 levels during *A. fumigatus* infection. (A) Representative flow cytometry plots and bar graphs (B-E) that indicate the frequency of RFP^+^ (CXCL9^+^; red bar in B-C), RFP^-^ (CXCL9^-^; gray bar in B-C), BFP^+^ (CXCL10^+^; blue bar, in D-E), and BFP^-^ (CXCL10^−^; gray bar in D-E) bystander and fungus-engaged leukocytes isolated from (A, left column) naïve or (A, middle and right column, B-E) infected Rex3 Tg → C57BL/6.SJL BM chimeric mice (n = 7) with 3 × 10^7^ AF633-labeled CEA10 conidia. The solid black gates in (A) indicate bystander neutrophils, while the solid blue gates indicate fungus-engaged leukocytes. The dashed line in the black and blue gate indicate CXCL9^+^ and CXCL9^-^ (top quadrants) leukocytes, and CXCL10^+^ and CXCL10^−^ (bottom quadrants) leukocytes. (F) Lung CXCL9 and (G) CXCL10 levels in naïve wild-type (WT, n = 5) and in WT (n = 10), *Clec7a*^-/-^(n = 8), *Card9*^-/-^ (n = 7), *Ifnar1*^-/-^ (n = 9), and *Ifnlr*^*-/-*^ (n = 9) mice 48 h pi with 3 × 10^7^ CEA10 conidia. Data were pooled from 2 independent experiments. (B-G) Dots represent individual mice and data are presented as mean ± SEM. Statistical analysis: Mann-Whitney test. See also Figure S2.

Dectin-1 is a major *A. fumigatus* recognition receptor by binding to *β*-glucan moieties that are exposed during conidial swelling, the first step in germination and hyphal formation (Brown et al., 2018; Hohl et al., 2005; Steele et al., 2005; Werner et al., 2009). Dectin-1 transduces signals via a cytoplasmic ITAM-like motif that, upon tyrosine phosphorylation, activates spleen tyrosine kinase and Card9 (Gross et al., 2006; Hsu et al., 2007a; Jhingran et al., 2015; Jia et al., 2014; Rogers et al., 2005). To determine whether Dectin-1/Card9 signaling mediates CXCL9 and CXCL10 production, we infected gene-deficient mice and found that CXCL9, but not CXCL10 production, was significantly attenuated in both *Clec7a*^-/-^ and *Card9*^-/-^ mice (Figures 2F-2G), directly linking fungal recognition by C-type lectin receptor signaling to CXCL9 production in the lung.

Prior studies have demonstrated that CXCL10 expression can be induced in a variety of cells, including endothelial cells, keratinocytes, fibroblasts, mesangial cells, astrocytes, monocytes, and neutrophils by stimulation with IFN-α, IFN-β, IFN-γ, or LPS and in T cells by antigen activation (Colvin et al., 2004; Luster and Ravetch, 1987; Ohmori and Hamilton, 1990; Qian et al., 2007). Since type I and type III interferon signaling are essential for host defense against *A. fumigatus* (Espinosa et al., 2017), we measured CXCL10 production in IFN-signaling deficient mice. Unlike CXCL9, lung CXCL10 levels were significantly reduced in *Ifnar1*^-/-^ and in *Ifnlr*^-/-^ mice (Figures 2F-2G), indicating that distinct upstream signals couple fungal recognition to CXCL9 and CXCL10 production during acute *A. fumigatus* infection.

Since *β*-glucan recognition and IFN signaling may impact the number of Mo-DCs and neutrophils in *A. fumigatus*-infected lungs (Jhingran et al., 2015; Jhingran et al., 2012; Werner et al., 2009), we sorted Mo-DCs and neutrophils from gene-deficient and wild-type mice and quantified *cxcl9* and *cxcl10* mRNA expression. Preliminary studies that measured *cxcl9* and *cxcl10* mRNA expression in Mo-DCs and neutrophils isolated from infected wild-type, *Clec7a*^*-/-*^, *Ifnar*^*-/-*^, and *Ifnlr*^*-/-*^ mice supported the concept that *β*-glucan recognition and IFN signaling directly regulate *cxcl9* and *cxcl10* transcription, respectively (data not shown).

### CXCR3 is critical for survival and fungal clearance

To assess the relevance of CXCR3 ligand production (Groom et al., 2012) for host outcomes, the survival of *Cxcr3*^-/-^ mice was compared to *Cxcr3*^+/+^ controls following *A. fumigatus* challenge. *Cxcr3*^-/-^ mice were significantly more susceptible to challenge compared to controls (Figure 3A). The heightened susceptibility correlated with an increase in lung fungal burden (Figures 3B, S3A and S3B) and in lung tissue damage, as determined by comparisons of bronchoalveolar lavage fluid (BALF) lactate dehydrogenase (LDH) (Figure 3C) and albumin (Figure 3D) levels at 48 h pi or 72 h pi. These data linked *A. fumigatus*-triggered CXCL9 and CXCL10 production by Mo-DCs and neutrophils during acute *A. fumigatus* challenge to CXCR3-dependent protection and fungal clearance.

**Figure 3.**
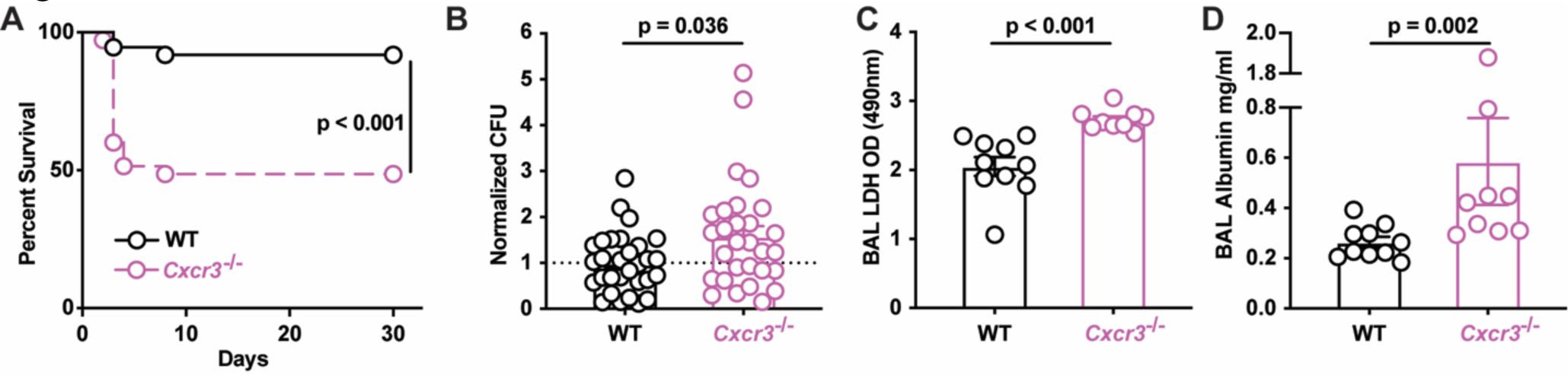
CXCR3 is critical for anti-*Aspergillus* defense. (A) Kaplan-Meier survival of C57BL/6 (n = 37) and *Cxcr3*^-/-^ (n = 35) mice challenged with 4-8 × 10^7^ CEA10 conidia. Data from 3 experiments were pooled. (B) Normalized lung fungal burden, (C) bronchoalveolar lavage fluid (BAL) LDH level, and (D) BAL albumin levels in C57BL/6 and *Cxcr3*^-/-^ mice 48 h pi with 3 × 10^7^ CEA10 conidia. (A-D) Dots represent individual mice and data were pooled from 2-4 experiments and presented as mean ± SEM. Statistical analysis: Mann-Whitney test. See also Figure S3.

### CXCR3- and CCR2-dependent signals mediate pDC lung recruitment

To define the cellular target of CXCR3 signaling, we measured CXCR3 expression on a wide range of leukocytes. Bone marrow (BM)-resident pDCs from naïve mice and lung pDCs from *A. fumigatus*-infected mice expressed CXCR3, unlike other myeloid cell subsets analyzed (Figure 4A). As expected, CD4^+^ T cells, CD8^+^ T cells and NK cells expressed CXCR3 (Figure S4A), but mice that lack the lymphoid lineage (*Rag2*^-/-^*Il2rg*^-/-^) did not exhibit heightened susceptibility to *A. fumigatus* challenge (Figure S4B), as reported previously (Espinosa et al., 2014).

**Figure 4.**
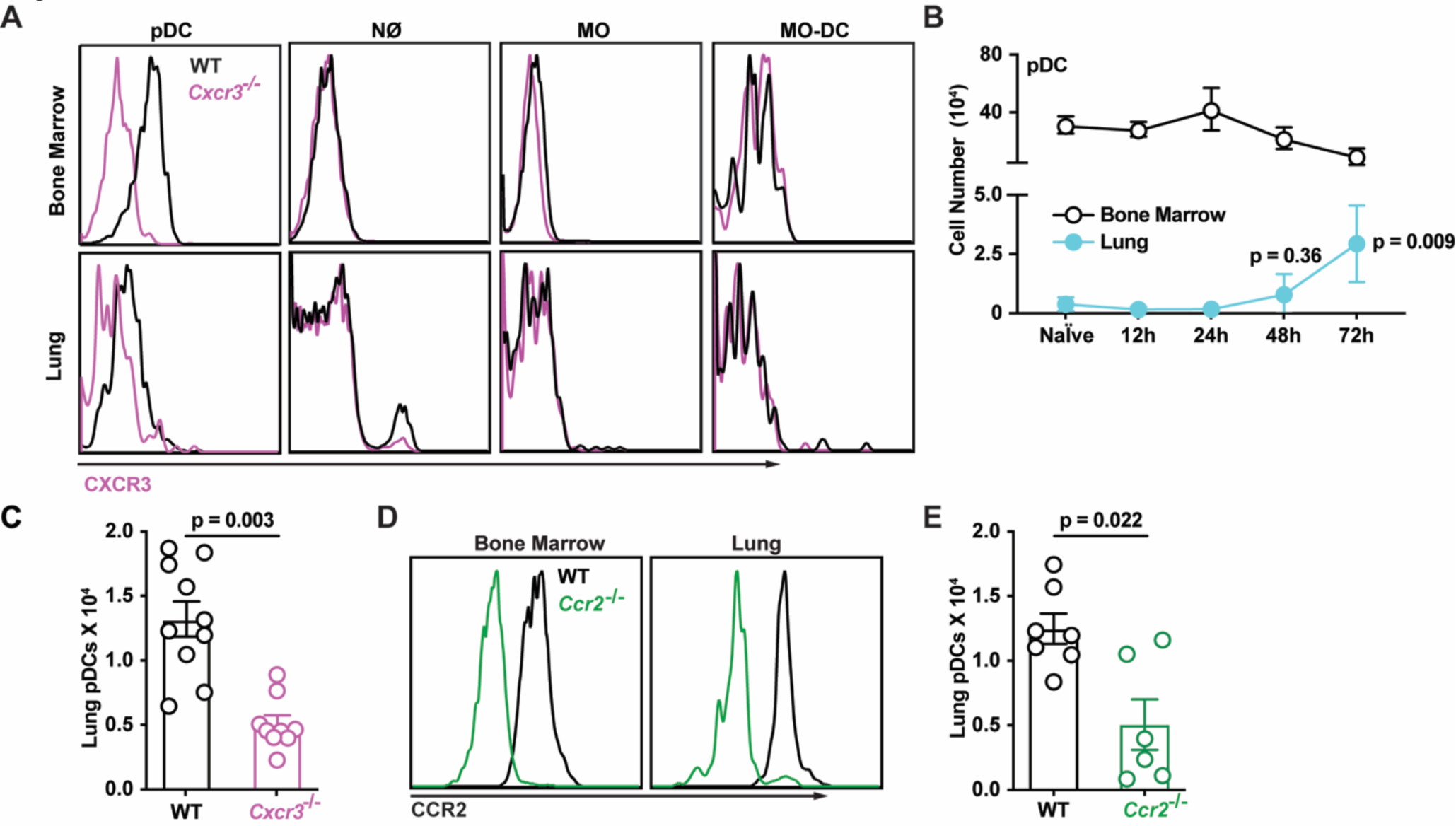
CXCR3^+^ pDCs traffic to *A. fumigatus*-infected lungs. (A) CXCR3 surface expression in BM (top row) and lung (bottom row) leukocytes isolated from WT (black lines) or *Cxcr3*^-/-^ mice (purple lines). (B) Lung (blue filled dots) and BM (open black dots) pDC numbers at baseline and indicated times pi with 3 × 10^7^ CEA10 conidia (n = 5). (C) Lung pDC numbers in WT (open black dots) and *Cxcr3*^-/-^ mice (open purple dots) at 72 h pi (n = 10). (D) CCR2 surface expression in BM and lung pDCs in WT (black lines) or *Ccr2*^-/-^ mice (green lines). (E) Lung pDC numbers in WT (open black dots) and *Ccr2*^-/-^ mice (open green dots) at 72 h pi (n = 10). (B, C, E) Data from 2-3 experiments were pooled and expressed as mean ± SEM. Statistical analysis: Mann-Whitney test. See also Figure S4.

On the basis of these results and published work (Ramirez-Ortiz et al., 2011), we hypothesized that CXCR3-dependent pDC trafficking is critical for innate antifungal immunity. In prior studies, we did not identify pDCs in lung cell suspensions due to the inclusion of collagenase type IV. This preparation method degraded a pDC epitope utilized for their identification (Figure S4C). Using a collagenase-free method to prepare single cell lung suspensions, we observed that pDCs started to infiltrate *A. fumigatus*-infected lungs at 48 h pi, with a peak at 72 h pi (Figure 4B). pDC lung infiltration followed the observed peak in lung CXCL9 and CXCL10 levels and the peak influx of CXCL9- and CXCL10-producing innate immune cells (Figures S4D-S4F). *Cxcr3*^-/-^ mice exhibited a ∼50% reduction in lung pDCs (Figure 4C), while the number of lung monocytes, Mo-DCs, and neutrophils were similar to control mice at 72 h pi (Figures S4G-S4I). CXCR3 surface expression was reduced on lung pDCs compared to circulating pDCs, consistent with the idea that CXCR3 surface expression is downregulated following engagement by CXCR3 ligands (Figure 4A). To examine whether CXCR3 acts in series or in parallel with other chemokine receptors to mediate pDC trafficking from the BM to the lung, we measured pDC surface expression of additional chemokine receptors in the resting and fungus-infected state. Consistent with prior studies (Fujimura et al., 2015; Sawai et al., 2013; Serbina and Pamer, 2006; Swiecki et al., 2017), we found that CCR2 is expressed by lung-infiltrating pDCs (Figure 4D). pDC lung recruitment was attenuated by 60-70% in *Ccr2*^-/-^ mice at 72 h pi compared to controls (Figure 4E). As expected (Hohl et al., 2009), lung monocyte and Mo-DC numbers were significantly reduced in *Ccr2*^-/-^ mice (Figures S4J and S4K), while lung neutrophil accumulation was not affected (Figure S4L). In sum, these data indicate that CXCR3 and CCR2 both mediate pDC trafficking.

### Sequential CCR2- and CXCR3-dependent signals mediate pDC lung recruitment

To determine at which steps CXCR3 and CCR2 regulate pDC trafficking, we generated mixed BM chimeric mice to compare the trafficking of congenically marked gene-deficient (*Cxcr3*^−/−^ or *Ccr2*^*-/-*^) and gene-sufficient (*Cxcr3*^+/+^ or *Ccr2*^*+/+*^) pDCs (Figure 5A). In one model, CXCR3 and CCR2 may act in parallel at the same trafficking step. In this scenario, the ratio of gene-knockout to gene-sufficient pDCs would be increased in the same compartment for both chemokine receptors, prior to the dependent trafficking step. In an alternate model, CXCR3 and CCR2 may act in series at different trafficking steps. In this scenario, the ratio of gene-deficient to gene-sufficient pDCs would be elevated in distinct compartments.

**Figure 5.**
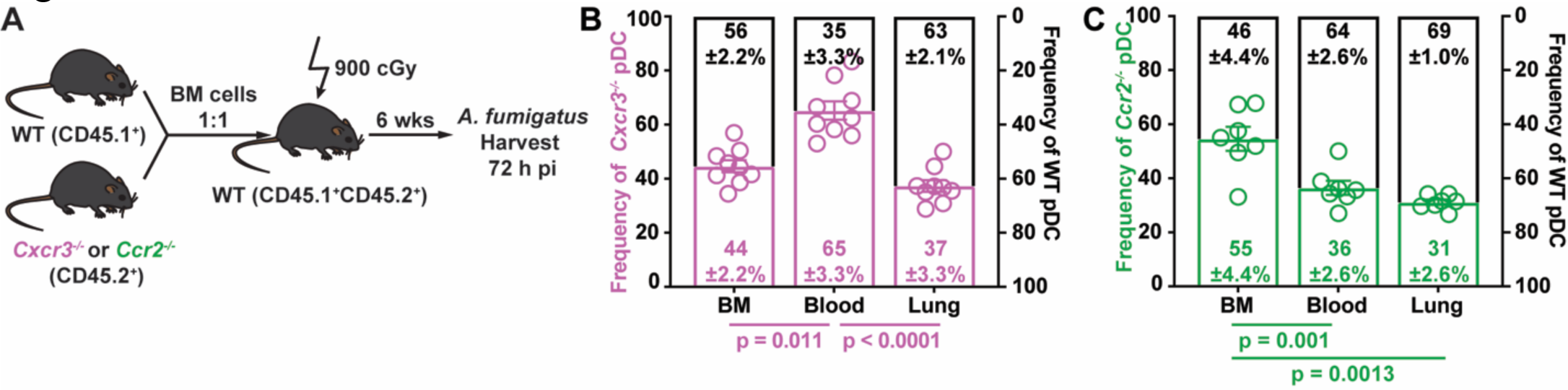
Sequential CCR2- and CXCR3-dependent signals mediate lung pDC recruitment. (A) Experimental scheme to generate mixed bone marrow chimeric mice and compare the trafficking of *Cxcr3*^*-/-*^, *Ccr2*^*-/-*^, and WT pDCs during *A. fumigatus* infection. (B, C) Relative frequencies of (B) *Cxcr3*^-/-^ (open purple bars) and *Cxcr3*^+/+^ (WT, open black bars) and (C) *Ccr2*^-/-^ (open green bars) and *Ccr2*^+/+^ (WT, open black bars) pDCs in the BM, blood, and lung of mixed BM chimeric mice 72 h pi. Data from 2-3 experiments were pooled and expressed as mean ± SEM. Statistical analysis: Mann-Whitney test. See also Figure S5.

The ratio of *Cxcr3*^−/−^ to *Cxcr3*^+/+^ pDCs was elevated in the blood compared to the BM and lung compartments (Figures 5B and S4A), both during infection and at baseline (Figure 5B and Figure S5A). These data support a model in which CXCR3 primarily, but not exclusively, mediates pDCs trafficking from the circulation into the lung in *A. fumigatus*-infected mice. In contrast to pDCs, the ratio *Cxcr3*^−/−^ to *Cxcr3*^+/+^ neutrophils and monocytes was similar in all compartments analyzed, consistent with CXCR3-independent trafficking (Figure S5B-S5C).

The ratio of *Ccr2*^−/−^ to *Ccr2*^+/+^ pDCs was elevated in BM compared to the blood and lung under all conditions examined. These data are consistent with a CCR2-dependent defect in pDC exit from the BM during infection (Figure 5C) and in the steady state (Figure S5D). As anticipated, monocyte bone marrow egress was highly dependent on CCR2 (Figure 5SE), while neutrophil bone marrow was not (Figure S5F). In sum, these data show that CCR2- and CXCR3-dependent signals act in series to mediate pDC entry into the circulation and pDC egress from the circulation into the lung, respectively.

### Lung pDCs are essential for host defense against *A. fumigatus*

To link CXCR3 signaling to pDC-dependent antifungal immunity, our model would predict that the chemokine receptor, CXCR3, and its cellular target, the pDC, are both essential for innate antifungal immunity. To test this conjecture, we infected BDCA2-DTR^+/-^ (pDC Depleter) mice in which pDCs are ablated at rest and under inflammatory conditions (Swiecki et al., 2010). Following DT treatment, pDCs were depleted fully in the steady state and at 72 h pi, while lung-infiltrating monocytes, Mo-DCs, neutrophils, and all other leukocyte subsets examined were not (Figures 6A-6E). In contrast to BDCA2-DTR^+/-^ mice, CCR2-DTR^+/-^ mice facilitated DT-induced ablation of pDCs in addition to other known CCR2^+^ cells, i.e. monocytes, Mo-DCs, and NK cells (Figures S6A-6E) (Espinosa et al., 2014; Hohl et al., 2009).

**Figure 6.**
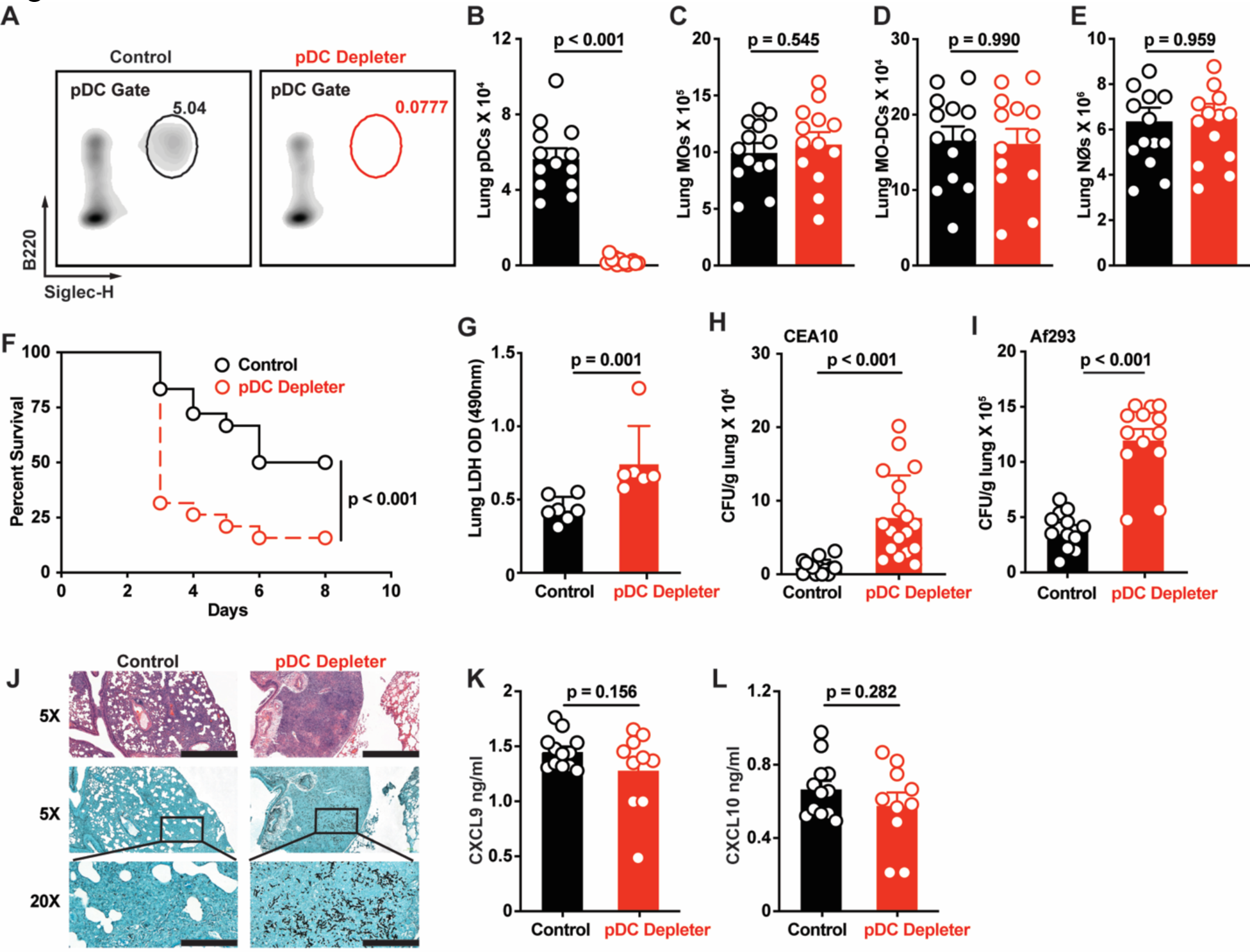
pDCs are critical for anti-*Aspergillus* defense. (A) Representative flow cytometry plots of B220^+^Siglec-H^+^ pDCs and (B) lung pDC, (C) lung monocyte, (D) lung Mo-DC, and (E) lung neutrophil numbers in DT-treated pDC Depleter mice (BDCA2-DTR; open red symbols) and non-Tg littermate controls (open black symbols) at 72 h pi with 3 × 10^7^ CEA10 conidia. (F) Kaplan Meier survival of DT-treated pDC Depleter (n = 20) and non-Tg littermate control mice (n = 19) infected with 3-6 × 10^7^ CEA10 conidia. (D) BAL LDH levels and (H) fungal burden of DT-treated pDC Depleter (open red symbols; n = 6-18) or non-Tg littermates (Control, open black symbols; n = 6-17) at 72 h pi with 3 × 10^7^ CEA10 conidia. (I) Lung fungal burden at 72 h pi with 3 × 10^7^ Af293 conidia. (K) Lung histopathology (top row, hematoxylin and eosin stain, scale bar 800μm; middle row, Gomori methenamine silver stain, scale bar 800μm; bottom row, GMS, scale bar 200μm) of DT-treated pDC Depleter (right column; n = 6) or non-Tg littermates (Control, left column; n = 6) at 72 h pi with 3 × 10^7^ CEA10 conidia. (K) Lung CXCL9 and (L) lung CXCL10 levels in DT-treated pDC Depleter mice (red symbols) and non-Tg littermate controls (black symbols) at 72 h pi with 3 × 10^7^ CEA10 conidia. (B-L) Data were pooled from 2-3 independent experiments and expressed as mean ± SEM. Statistical analysis: Mann-Whitney test. See also Figure S6.

To define the functional role of pDCs during *A. fumigatus* infection, we compared the mortality, lung LDH levels, lung fungal burden and histopathology in pDC-depleted mice and in littermate controls. pDC-depleted mice were more susceptible to *A. fumigatus* challenge than control mice (Figure 6F) and mortality correlated with greater lung damage, as measured by BAL fluid lactate dehydrogenase (LDH) levels (Figure 6G), an increased fungal burden following infection with two commonly used *A. fumigatus* isolates (Figures 6H and 6I), and a failure to control fungal germination in the lung at 72 h pi (Figure 6J). Importantly, pDCs did not contribute to the production of CXCL9 and CXCL10 in the lung during *A. fumigatus* challenge, as predicted by prior data (Figures 6K and 6L). In sum, CXCR3^+^ pDCs were essential for innate antifungal immunity in the lung after *A. fumigatus* challenge.

### Lung pDCs enhance neutrophil fungicidal activity

To determine how pDCs shape innate antifungal immunity in the lung, we hypothesized that pDCs are critical for antifungal effector functions, since lung neutrophil and Mo-DC numbers were unaffected in pDC depleted mice. To test this hypothesis, we utilized the fluorescent *Aspergillus* reporter (FLARE) conidia that encode a dsRed viability fluorophore and are labeled with an Alexa Fluor 633 tracer (AF633). FLARE conidia enable us to distinguish live (DsRed^+^AF633^+^) and dead (DsRed^-^AF633^+^) conidia during cellular interactions with leukocytes with single-encounter resolution (Figure 7A). Using the FLARE strain, we quantified leukocyte conidial uptake and killing in pDC-depleted and littermate control mice. pDCs did not bind to or engulf *A. fumigatus* conidia, as measured by acquisition of AF633 fluorescence (Figures 7B and S7A). pDC ablation did not affect lung neutrophil fungal uptake at 72 h pi compared to lung neutrophils in pDC-sufficient controls (Figures 7C and 7D). However, the frequency of neutrophils that contained live conidia was substantially increased in pDC-depleted mice compared to control mice (Figures 7C and 7E). In other words, neutrophil-engulfed conidia were more likely to be killed in control mice than in pDC-depleted mice (Figures 7C and 7E). pDC ablation did not alter conidial uptake or significantly reduce conidial killing by monocytes and Mo-DCs at 72 h pi (Figures 7D and 7E). These findings indicate that pDCs enhanced neutrophil fungicidal activity.

**Figure 7.**
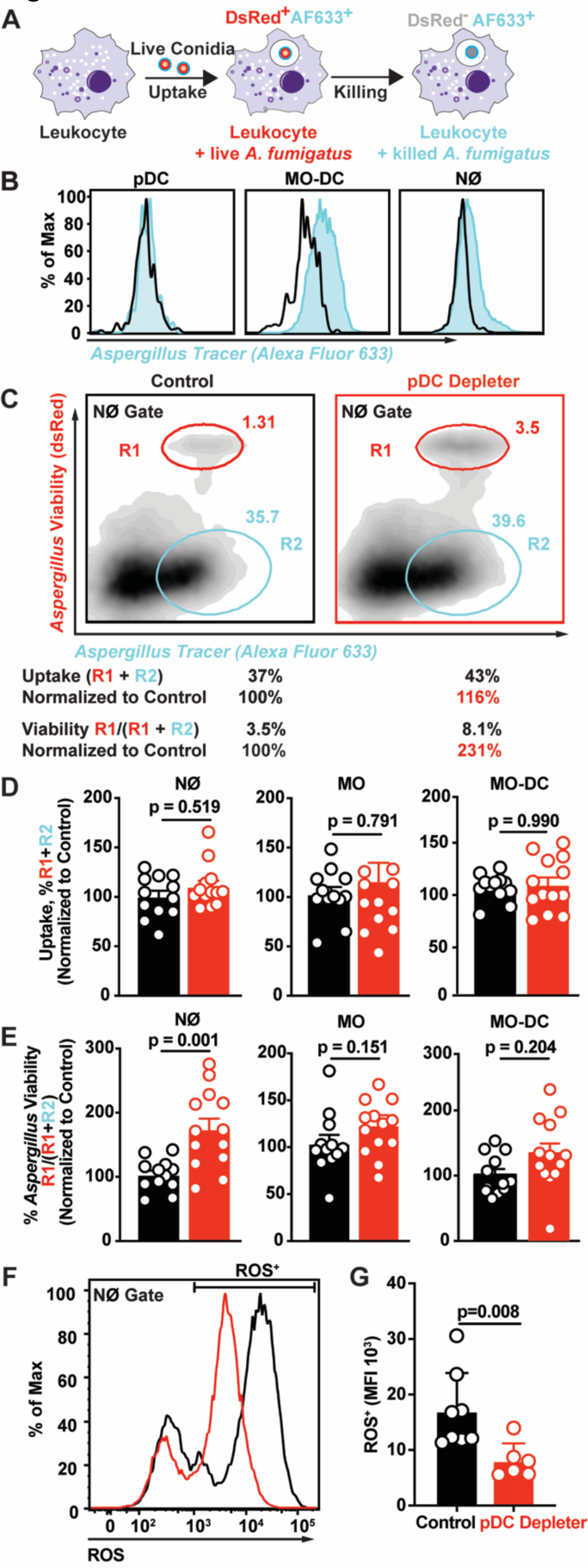
Lung pDCs enhance neutrophil fungicidal activity. (A) Schematic of FLARE strain and changes in fluorescence emission following fungal uptake and killing by host leukocytes. (B) AF633 fluorescence intensity in indicated lung leukocytes 72 h pi with FLARE (blue line) or AF633-unlabeled conidia. (C) Representative plots that display dsRed and AF633 fluorescence intensity of lung neutrophils in DT-treated pDC Depleter (right panel) and non-Tg littermates (left panel) 72 h pi with 3 × 10^7^ Af293 FLARE conidia. R1 denotes neutrophils that contain live conidia, R2 denotes neutrophils that contain killed conidia. (D and E) The plots show neutrophil, monocyte, and Mo-DC (D) conidial uptake (R1 + R2) ± SEM and (E) conidial viability (R1/ (R1 + R2) ± SEM in indicated lung leukocytes isolated from DT-treated pDC Depleter (red symbols) and non-Tg littermates (black symbols) 72 h pi with 3 × 10^7^ FLARE conidia. (F) Representative plot and (G) mean ± SEM neutrophil ROS production in cells isolated from DT-treated pDC Depleter (right panel) and non-Tg littermates (left panel) 72 h pi with 3 × 10^7^ CEA10 conidia. (C-E) Data from 2 experiments were pooled. (F, G) Data are representative of 2 experiments. (D, E, G) Dots represent individual mice and data are expressed as mean ± SEM. Statistical analysis: Mann-Whitney test. See also Figure S7.

Neutrophils generate reactive oxygen species (ROS) via NADPH oxidase as a primary effector mechanism against *A. fumigatus* (Marciano et al., 2015). Neutrophil NADPH oxidase induces a regulated cell death process in *A. fumigatus* conidia (Shlezinger et al., 2017). To determine whether pDCs regulate the neutrophil oxidative burst in *A. fumigatus*-infected mice, we measured neutrophil ROS production in pDC-depleted mice. The ROS median fluorescence intensity in ROS^+^ lung neutrophils isolated from pDC-depleted mice was significantly reduced compared to ROS^+^ neutrophils isolated from littermate control mice at 72 h pi (Figure 7E and 7F). These data show that lung pDCs regulate neutrophil ROS generation during *A. fumigatus* challenge.

## Discussion

Humans maintain lifelong sterilizing immunity to inhaled mold conidia. Breaches in the respiratory innate immune system lead to mold pneumonia, a life-threatening disease in patients with functional or numeric deficits in myeloid cell function, particularly with the loss of the antifungal properties of neutrophils. In this study, we demonstrate a three-cell circuit that involves pDCs, monocytes and derivative Mo-DCs, and neutrophils to regulate sterilizing immunity against *Aspergillus* conidia. This circuit acts as a feedforward amplification mechanism, by coupling fungal recognition and fungus-induced inflammation to CXCR3 signaling-dependent pDC recruitment in the lung. pDCs regulate neutrophil ROS induction and fungal killing to mediate sterilizing immunity at the respiratory barrier.

Our data demonstrate that conidial uptake by lung-infiltrating monocytes and Mo-DCs triggered Dectin-1- and CARD9-dependent production of CXCL9 and type I and type III interferon signaling-dependent production of CXCL10; the latter process was readily detectable in neutrophils as well. The relative contribution of CXCL9 and CXCL10 and potential molecular redundancy with regard to downstream CXCR3-dependent processes during fungal infection remains an open question for future studies. Prior studies identified a polymorphism in the *Cxcl10* gene as a risk factor for IA in allogeneic hematopoietic cell transplant patients (Fisher et al., 2017; Mezger et al., 2008). In this cohort, patients with IA had higher CXCL10 serum levels than controls that did not develop IA, consistent with the idea that pulmonary fungal infection induces CXCL10, similar to findings in mice in this work. Furthermore, the high-risk haplotype was associated with reduced *Cxcl10* mRNA expression by immature DCs (iDCs) co-cultured with *Aspergillus* germlings compared to wild-type iDCs (Mezger et al., 2008). Thus, human iDCs and murine lung Mo-DCs both responded to *A. fumigatus* challenge by transcribing *Cxcl10* mRNA. Although this process is generally regulated by IFN signaling in both mammalian species (O’Connell et al., 2019; Ohmori et al., 1993), a recent study by Rivera and colleagues defined essential roles for type I and type III, but not type II, IFNs in murine host defense against *A. fumigatus* (Espinosa et al., 2017). One model to account for these findings is that an early wave of type I interferon signaling, detected as early as 6 h pi in the lung and produced primarily by CCR2^+^ lung-infiltrating monocytes and Mo-DCs (Espinosa et al., 2017), may drive CXCL10 release in the lung in a paracrine or autocrine manner, as observed in this study. In accordance with this model, CXCL10 lung levels that peak at 48 h pi are dependent on type I and type III IFN receptor signal transduction in the *Aspergillus*-infected murine lung.

Although IFN-dependent and -independent stimuli can induce CXCL9 during microbial infection (Forero et al., 2019; Groom and Luster, 2011), previous studies have not directly linked C-type lectin receptor signaling to fungus-induced CXCL9 production. In murine models of experimental cryptococcosis with a wild-type or a recombinant vaccine strain that expresses human IFN-*γ* fungal infection led to lung CXCL9 and CXCL10 production, though the identity of chemokine-producing cells was not investigated (Hole et al., 2016; Yamamoto et al., 2014). In this model, *Card9*^-/-^ mice had lower lung CXCL9 levels than control mice. This finding was attributed to reduced lung infiltration of IFN-*γ*-producing NK and CD4^+^ T cells because *Cryptococcus*-stimulated BM-DCs failed to produce either cytokine *in vitro* (Yamamoto et al., 2014). In contrast, splenic and central nervous system (CNS)-resident Mo-DCs produced copious levels of CXCL9 and CXCL10 during murine cerebral malaria (Hirako et al., 2016). Because a significant fraction of CXCL9^+^CXCL10^+^ splenic Mo-DCs contained hemozoin, a malarial pigment, this study supported the idea that microbial uptake regulated CXCR3 ligand release either directly or indirectly in infected cells. The authors provided evidence for indirect CXCL9 and CXCL10 regulation since their secretion was highly dependent on intact IFN-*γ* receptor signaling. In the case of respiratory *A. fumigatus* infection, the dual regulation of CXCL9 by Dectin-1/Card9 and CXCL10 by type I and type III IFN signaling indicates both direct and indirect regulation of distinct CXCR3 ligands by a pathogen-associated molecular pattern receptor and soluble type I and type III IFNs.

CXCR3 ligands have been studied extensively in the context of CD4^+^ and CD8^+^ T cell trafficking in peripheral lymphoid and non-lymphoid tissues, in part due to high CXCR3 expression on effector and memory T cell subsets, in contrast to naïve T cells (Qin et al., 2011). In draining LNs, antigen-activated and transferred DCs express CXCL9 and CXCL10 and thereby regulate the frequency of IFN-*γ*^+^ CD4^+^ T cells, linking the CXCR3 signaling axis to the formation of effector Th1 cells (Groom et al., 2012). CXCR3 facilitated the entry of effector T cells into otherwise restricted sites, exemplified by CXCR3^+^ CD8^+^ T cells infiltration of the CNS or genital tract during murine malaria, West Nile virus, or herpes simplex virus 2 infection (Hirako et al., 2016; Nakanishi et al., 2009; Thapa and Carr, 2009; Thapa et al., 2008). During bacterial infection, a study reported that CXCR3 signaling regulated neutrophil influx to the infected cecum and thus controlled *Salmonella* extra-intestinal dissemination (Chami et al., 2017). In rodent models of viral (e.g., influenza virus or coronavirus) or chemical acute respiratory distress syndrome, CXCL10^+^ lung-infiltrating neutrophils and CXCL9^+^ macrophages mediated the recruitment of tissue-destructive CXCR3^+^ neutrophils; this pathologic process was ameliorated in *Cxcr3*^-/-^ or *Cxcl10*^-/-^ mice (Ichikawa et al., 2013). During respiratory fungal challenge, we did not observe CXCR3 expression on lung-infiltrating neutrophils and neutrophil recruitment was unaffected in *Cxcr3*^-/-^ mice compared to control mice. In more recent work in the *Salmonella* model, infected splenic macrophages reside in T cell-sparse granulomas but are surrounded by mononuclear phagocytes that release CXCL9 and CXCL10 to recruit antigen-specific CXCR3^+^ Th1 cells for bacterial containment (Goldberg et al., 2018). Thus, CXCR3 ligands target both innate and adaptive immune cells in a highly context-dependent manner; this process can result in microbial containment or microbe-elicited tissue damage.

pDCs originate in the bone marrow (BM), circulate in the blood, and traffic to lymphoid and nonlymphoid tissues under homeostatic and inflammatory conditions (Fujimura et al., 2015; Sozzani et al., 2010; Swiecki et al., 2017). However, the mechanism that couples microbial recognition to pDC accumulation at peripheral sites has remained poorly understood. During respiratory fungal infection, neither CCR2^+^ monocytes nor CCR2^-^ neutrophils had the capacity to respond to CXCR3 ligands because these cells do not express CXCR3. Thus, the recruitment of lung-infiltrating monocytes and pDCs occurs via a pathway that bifurcated after the initial CCR2-dependent egress from the bone marrow reservoir. Although the precise cues that regulate the CXCR3-independent entry of circulating monocytes into the fungus-infected lung remain unknown, integrin-mediated mechanisms enable monocytic influx into hepatic foci of *Listeria monocytogenes* during systemic infection (Shi et al., 2010).

During other types of lung infections, e.g. influenza and RSV, pDCs traffic to the lungs and draining LNs of mice (Langlois and Legge, 2010; Smit et al., 2006; Wang et al., 2006) via pathways that are distinct compared to conventional DCs (Sawai et al., 2013; Sozzani et al., 2010). For example, pDC LN trafficking during murine influenza is regulated primarily by L-selectin (CD62L), with contributions from CXCR3 and CXCR4. Further pDC positioning within the LN is mediated by CCR7 and the release of pDC-derived CXCR3 ligands to enable the formation of clusters of pDCs that interact and activate cells of the adaptive immune system (Krug et al., 2002). In contrast, we did not observe pDC-dependent release of CXCL9 and CXCL10 in the *Aspergillus*-infected lung, consistent with the model that pDC recruitment cues depends on heterologous myeloid cells. Beyond CXCR3, L-selectin, and CXCR4, the E-selectin ligands, *β*1 and *β*2 integrins, CCR5, CCR7, and the chemerin receptor ChemR23 have been implicated in pDC trafficking during states of inflammation (Diacovo et al., 2005; Kohrgruber et al., 2004; Sozzani et al., 2010; Vanbervliet et al., 2003; Wendland et al., 2007), though it has remained unclear how individual cues cooperate to guide distinct pDC trafficking steps, and what the relative strength of individual cues is for defined tissue destinations and types of inflammatory stimuli. The sequential requirements for CCR2 and CXCR3 signaling in distinct pDC trafficking steps during respiratory *A. fumigatus* infection provides a combinatorial model, in which a more general cue for myeloid cell bone marrow egress (i.e., CCR2 that acts on monocytes and pDCs) is combined with a more specific, pDC-targeted cue (i.e., CXCR3). The latter cue emanates primarily from CCR2^+^ Mo-DCs that were not equipped to control fungal tissue invasion or license the full spectrum of neutrophil antifungal activity per se.

pDCs were functionally characterized as mediators of antiviral defense (Assil et al., 2019; Swiecki and Colonna, 2010). Recent studies have expanded their role to anti-bacterial (*e*.*g*., *Legionella pneumophila, Chlamydia pneumoniae, Staphylococcus aureus*) (Ang et al., 2010; Crother et al., 2012; Parcina et al., 2013) and to anti-fungal defense (Ang et al., 2010; Hole et al., 2016). pDC-mediated antiviral responses are stimulated upon physical contact with infected cells (Assil et al., 2019). pDCs adhere to infected cells via α_L_β_2_ integrin/ICAM-1 interactions, during which viral RNA is transferred to pDCs, leading to IFN production via the nucleic acid sensor TLR7. This process activates type I IFN-dependent antiviral programs in infected tissues. During respiratory fungal infection, lung-infiltrating pDCs do not interact directly with *Aspergillus* conidia, but instead regulate neutrophils ROS induction. Our study does not exclude the possibility that *Aspergillus*- or host cell-derived nucleic acids activate pDCs in the lung in situ. Prior work has demonstrated that *Aspergillus*-derived unmethylated CpG sequences can activate TLR9 signaling *in vitro* (Herbst et al., 2015; Ramirez-Ortiz et al., 2008). Alternatively, pDCs may recognize fungal Dectin-2/CLEC4N or Dectin-3/CLEC4E ligands (Loures et al., 2015; Maldonado and Fitzgerald-Bocarsly, 2017; Preite et al., 2018). In patients with systemic lupus erythematosus, neutrophil-derived extracellular traps (NETs) activate pDCs by releasing DNA-protein complexes, resulting in TLR9-dependent release of pathologic type I IFNs during autoimmunity (Garcia-Romo et al., 2011; Lande et al., 2011). While *A. fumigatus* conidia are poor inducers of NETosis, products of germination, specifically hyphae that are too large to be phagocytosed, are potent stimuli for NETosis (Bianchi et al., 2009; Bruns et al., 2010; Gibrat et al., 1989). Thus, pDC activation in the fungus-infected lung likely occurs by multiple, yet undefined mechanisms.

The molecular basis for pDC-neutrophil crosstalk represents an important area of future investigation. Two non-exclusive models are likely operative. First, pDCs may form direct contact with neutrophils after arriving at the site of infection. In this model, an individual pDC would interact with many neutrophils since neutrophils are ∼100 times more prevalent in the lung at 48 h pi compared to pDCs. The relative paucity of pDCs, compared to other infiltrating myeloid cells (i.e., neutrophils, Mo-DCs), makes it unlikely that direct pDC-mediated fungistasis, observed at high effector to target cell ratios in the test tube, is physiologically significant in the fungus-infected lung (Loures et al., 2015; Maldonado and Fitzgerald-Bocarsly, 2017). Second, either fungus-triggered or host cell-triggered pDC activation may result in the release of soluble mediators that boost neutrophil fungicidal activity. Previous studies have identified GM-CSF, type I IFN, and type III IFN as important regulators of neutrophil NADPH oxidase activity during respiratory *A. fumigatus* and *B. dermatiditis* challenge (Espinosa et al., 2017; Hernandez-Santos et al., 2018; Kasahara et al., 2016). The development of pDC-restricted Cre-lox candidate gene deletion strategies is essential to test these candidates formally.

In this study, we uncover essential functions for pDCs in innate antifungal defense that cannot be compensated by the presence of other myeloid cells, including neutrophils, monocytes, and Mo-DCs. In fact, fungus-engaged Mo-DCs and neutrophils harness pDCs as a feedforward amplification mechanism to enhance innate antifungal activity in the lung by coupling fungal recognition and fungus-induced inflammation to the CXCR3 signaling-dependent recruitment of pDCs into the fungus-infected lung. In the lung pDCs regulate neutrophil ROS induction, a process that induces a regulated cell death process in mold conidia (Shlezinger et al., 2017). Our findings indicate that pDC recovery following administration of chemotherapy and in bone marrow transplant recipients may represent an important variable that affects infectious susceptibility not just to viral, but also to fungal pathogens. Recent studies in bone marrow transplant patients indicate that donor pDC reconstitution in the recipient is associated with favorable clinical outcomes, including non-relapse mortality from unrelated donors (Goncalves et al., 2015; Waller et al., 2019; Waller et al., 2014). These data raise the possibility that pDC recovery in bone marrow transplant patients may be important for controlling systemic fungal infections. Further studies to decipher the mechanistic role of pDCs in antifungal immunity is likely to inform strategies to harness these cells for prophylactic or therapeutic gain in vulnerable patient populations.

## ACKNOWLEDGMENTS

We thank Eric Pamer (University of Chicago) for sharing *Ccr2*^*-*/-^ and *Ifnar1*^-/-^ mice, Xin Lin (Tsinghua University) for *Card9*^*-/-*^ mice, Robert A. Cramer (Dartmouth College) for sharing the *A*.*fumigatus* CEA10 strain and for numerous conversations and insights into this study. We thank Franck Barrat, and Iliyan Iliev (both Weill Cornell Medical College), and members of the Hohl laboratory for insightful discussions. The studies were supported by Burroughs Wellcome Fund Investigator in the Pathogenesis of Infectious Diseases Awards (TMH and AR) and by NIH grants P30 CA 008748 (to MSKCC), R01 AI 093808 (TMH), R01 AI 139632 (TMH), R01 CA 204028 (ADL), R01 AI 114747 (AR), and R01 AI141368 (AR). The funders had no role in study design, data collection and analysis, decision to publish or preparation of manuscript.

## AUTHOR CONTRIBUTIONS

Conceptualization, T.M.H., Y.G. and S.K.; Methodology, T.M.H., Y.G. and S.K.; Investigation, Y.G., S.K. A.J., B.Z., M.A.A., K.A.M.M, V.E., and A.R.; Writing – Original Draft, Y.G. and T.M.H.; Writing – Review & Editing, Y.G. and T.M.H.; Funding Acquisition, T.M.H.; Resources, A.D.L., V.E., and A.R.

## STAR⋆METHODS

### KEY RESOURCES TABLE

| REAGENT or RESOURCE | SOURCE | IDENTIFIER |
| --- | --- | --- |
| Antibodies |  |  |
| APC Mouse anti-Mouse CD45.1 (clone A20) | BD Bioscience | Cat#558701; RRID: AB_1645214 |
| PerCP-Cy<sup>TM</sup>5.5 Mouse anti-Mouse CD45.2 (clone 104) | BD Bioscience | Cat#552950; RRID: AB_394528 |
| APC-Cy<sup>TM</sup>7 Rat anti-Mouse CD45 (clone 30-F11) | BD Bioscience | Cat#557659; RRID: AB_396774 |
| PerCP-Cy<sup>TM</sup>5.5 Rat anti-CD11b (clone M1/70) | BD Bioscience | Cat#550993; RRID: AB_394002 |
| PE-Cy<sup>TM</sup>7 Hamster anti-Mouse CD11c (clone HL3) | BD Bioscience | Cat#558079; RRID: AB_647251 |
| BV 605<sup>TM</sup> anti-mouse CD11c Antibody (clone N418) | BioLegend | Cat#117334; RRID: AB_117334 |
| PE-Cy<sup>TM</sup>7 Rat Anti-Mouse Ly-6C (clone AL-21) | BD Bioscience | Cat#560593; RRID: AB_1727557 |
| FITC Rat anti-Mouse Ly-6G (clone 1A8) | BD Bioscience | Cat#551460; RRID: AB_394207 |
| FITC Rat anti-Mouse CD45R/B220 (clone RA3-6B2) | BD Bioscience | Cat#553087; RRID: AB_394617 |
| PE anti-mouse CD183 Antibody (clone CXCR3-173) | BioLegend | Cat#126505; RRID: AB_1027656 |
| BV421 Rat Anti-Mouse Siglec-F (clone E50-2440) | BD Bioscience | Cat#562681; RRID: AB_2722581 |
| BV605 Rat Anti-Mouse Siglec-H (clone 440c) | BD Bioscience | Cat#747673; RRID: AB_2744234 |
| Purified Rat Anti-Mouse CD16/CD32 (clone 2.4G2) | BD Bioscience | Cat#553142; RRID: AB_394657 |
| eFluor 450 NK1.1 Monoclonal Antibody (clone PK136) | eBioscience | Cat#48-5941-80; RRID: AB_2043878 |
| eFluor 450 CD19 Monoclonal Antibody (clone eBio1D3) | eBioscience | Cat#48-0193-82; RRID: AB_2734905 |
| Alexa Fluor 700, I-A/I-E Monoclonal Antibody (clone M5/114.15.2) | eBioscience | Cat#56-53-21; RRID: AB_494009 |
| FITC Mouse IgG1 kappa Isotype Control | eBioscience | Cat#11-4714-42; RRID: AB_10596964 |
| PE Rat IgG2a kappa Isotype Control | eBioscience | Cat#12-4321-42; RRID: AB_1518773 |
| PerCP-Cyanine5.5 IgG Isotype Control | eBioscience | Cat#45-4888-80; RRID: AB_906260 |
| APC Rat IgG2b kappa Isotype Control | eBioscience | Cat#17-4031-82; RRID: AB_470176 |
| APC-eFluor 780 Rat IgG2b kappa Isotype Control | eBioscience | Cat#47-4031-80; RRID: AB_1272021 |
| Alexa Fluor 700 Rat IgG2b kappa Isotype Control | eBioscience | Cat#56-4031-80; RRID: AB_837123 |
| PE-Cyanine7 Rat IgG2a kappa Isotype Control | eBioscience | Cat#25-4321-82; RRID: AB_470200 |
| eFluor 450 Rat IgG1 kappa Isotype Control | eBioscience | Cat#48-4301-82; RRID: AB_1271984 |
| Cat#48-4301-82; RRID: AB_1271984 | BioLegend | Cat#400649; RRID: N/A |
| Bacterial and Virus Strains |  |  |
| AF293 | Fungal genetics stock center | #A1100 |
| AF293 ds-Red | (Jhingran et al., 2012) | N/A |
| CEA10 (also known as CBS144.89) | (Girardin et al., 1993) | Received from Robert A. Cramer |
| Chemicals, Peptides, and Recombinant Proteins |  |  |
| Alexa Fluor 633 succinimidyl ester | Invitrogen | Cat#S12375 |
| Collagenase IV | ThermoFisher Scientific | Cat#LS004189 |
| RPMI-1640 | RPMI-1640 | 21870092 |
| Enrofloxacin | Bayer | Baytril 100 |
| Voriconazole | Pfizer | N/A |
| Tween-20 | Sigma | P9416 |
| Protease Inhibitor Cocktail | Roche | Cat#11697498001 |
| DNase I, Grade II | Sigma | Cat#10104159001 |
| BCG albumin assay kit | Sigma | MAK124 |
| isoflurane | Henry Schein Animal Health | Cat#29405 |
| Diphtheria toxin (DT) | List Biological Laboratories | Cat#150 |
| Paraformaldehyde, 32% Solution | ThermoFisher Scientific | Cat#50-980-495 |
| CM-H<sub>2</sub>DCFDA | ThermoFisher Scientific | Cat#C6827 |
| 10× HBSS | ThermoFisher Scientific | 14065056 |
| TRIzol-LS | Invitrogen | 10296028 |
| Mouse CXCL9/MIG DuoSet ELISA | R&D systems | DY492 |
| Mouse CXCL10/IP-10/CRG-2 DuoSet ELISA | R&D systems | DY466 |
| CytoTox96 non-radioactive cytotoxicity assay kit | Promega | G1780 |
| High Capacity RNA-to-cDNA Kit | Appliedbiosystems | 4387406 |
| Ribosomal RNA Control Reagents | Appliedbiosystems | 4308329 |
| *Cxcl9* ASSAY ID: Mm01345159_m1 | Thermo Fisher | 4448892 |
| *Cxcl10* ASSAY ID: Mm00445235_m1 | Thermo Fisher | 4453320 |
| Experimental Models: Organisms/Strains |  |  |
| *Cxcr3*<sup>-/-</sup> | The Jackson Laboratory | JAX: 005796 |
| BDCA2-DTR | The Jackson Laboratory | JAX: 014176 |
| *Clec7a*<sup>-/-</sup> | (Saijo et al., 2007) | N/A |
| *Card9*<sup>-/-</sup> | (Hsu et al., 2007) | JAX:028652 |
| *Ccr2*<sup>-/-</sup> | The Jackson Laboratory | JAX: 004999 |
| *Ifnar1*<sup>-/-</sup> | The Jackson Laboratory | JAX: 32045 |
| *Ifnlr*<sup>-/-</sup> | (Lin et al., 2016) | N/A |
| C57BL/6 | The Jackson Laboratory | JAX: 000664 |
| C57BL/6.SJL | Charles River Laboratories | Stock No. 564 |
| *Rag2*<sup>-/-</sup>*Il2rg*<sup>-/-</sup> | Taconic | JAX: 4111 |
| MRP8-Cre-IRES/GFP | The Jackson Laboratory | JAX: 021614 |
| Oct-4/rtTA | The Jackson Laboratory | Oct-4/rtTA |
| CCR2-DTR | (Hohl et al., 2009) | N/A |
| REX3 Tg | (Groom et al., 2012) | N/A |
| Software and Algorithms |  |  |
| Prism 8 | Prism 8 | N/A |
| Flow Jo 9.9.6 | Flow Jo 9.9.6 | N/A |

## RESOURCE AVAILABILITY

Further information and requests for resources or reagents should be directed to the Lead Contact, Tobias M. Hohl.

### Materials Availability

This study did not generate new unique reagents.

### Data and Code Availability

This study did not generate or analyze new datasets or codes.

## EXPERIMENTAL MODEL AND SUBJECT DETAILS

### Mice

*Cxcr3*^-/-^, C57BL/6 (CD45.2^+/+^), BDCA2-DTR mice were purchased from Jackson Laboratories (JAX: 014176). *Rag2*^*-/-*^*Il2rg*^-/-^ mice (Stock No. 4111) were purchased from Taconic. *C*cr2^-/-^ (JAX: 004999) (Serbina and Pamer, 2006), *Ifnar1*^-/-^ (JAX: 32045) (Jia et al., 2009), and *Ifnlr*^*-/-*^ (Lin et al., 2016) mice were provided by Dr. Eric Pamer (University of Chicago) and Dr. Sergei Kotenko (Rutgers), respectively. Bones from REX3 Tg (CXCL9-Red Red fluorescent protein/CXCL10-Blue fluorescent protein reporter) mice (Groom et al., 2012) were provided by Dr. Andrew Luster (Massachusetts General Hospital). *Clec7a*^-/ -^ mice (Saijo et al., 2007) were provided by Dr. Shinobu Saijo (University of Tokyo). *Card9*^-/-^ mice (Hsu et al., 2007b) were provided by Dr. Xin Lin (Tsinghua University). MRP8-Cre-IRES/GFP and Oct-4/rtTA mice were crossed to generate ROSA26-iDTR^Mrp8-Cre^ mice. C57BL/6 and C57BL/6.SJL mice were crossed to generate CD45.1^+^CD45.2^+^ recipient mice for mixed BM chimeras. CCR2-DTR^+/-^ (Hohl et al., 2009) and non-transgenic littermates on the *Rag2*^-/-^*Il2rg*^-/-^ background were maintained on amoxicillin- and Vitamin E-containing chow. All experiments with CCR2-DTR, CCR2-DTR *Rag2*^-/-^*Il2rg*^-/-^ and BDCA2-DTR mice used co-housed littermate controls. For experiments in which the breeding strategy did not yield littermate controls, gene-knockout mice were co-housed with C57BL/6 mice for 14 days prior to infection, whenever possible. All mouse strains were bred and housed in the MSKCC or Rutgers Research Animal Resource Center under specific pathogen-free conditions. All animal experiments were conducted with sex- and age-matched mice and performed with MSKCC or Rutgers Institutional Animal Care and Use Committee approval. Animal studies complied with all applicable provisions established by the Animal Welfare Act and the Public Health Services Policy on the Humane Care and Use of Laboratory Animals.

### Generation of Bone Marrow Chimeric Mice

For single BM chimeras, CD45.1^+^ C57BL/6.SJL recipients were lethally irradiated (900cG), reconstituted with either 2-5 × 10^6^ CD45.2^+^ Rex3-Tg, CD45.2^+^ ROSA26-iDTR^Mrp8-Cre^, or CD45.2^+^ non-Cre iDTR littermates BM cells. For mixed BM chimeras, CD45.1^+^CD45.2^+^ recipients were irradiated and reconstituted with a 1:1 mixture of CD45.1^+^ C57BL/6.SJL and CD45.2^+^ *Cxcr3*^-/-^ or CD45.2^+^ *Ccr2*^-/-^ BM cells. After BM transplantation, recipient mice received 400 μg/ml enrofloxacin in the drinking water for 21 days to prevent bacterial infections and rested for 6-8 weeks prior to experimental use.

### *Aspergillus fumigatus* culture and Infection Model

*A. fumigatus* Af293, Af293-dsRed (Jhingran et al., 2012), and CEA10 (Girardin et al., 1993) strains were cultured on glucose minimal medium slants at 37°C for 4-7 days prior to harvesting conidia for experimental use. To generate FLARE conidia, briefly, 7 × 10^8^ Af293-dsRed conidia were rotated in 10 μg/ml Biotin XX, SSE in 1 ml of 50 mM carbonate buffer (pH 8.3) for 2 hr at 4 ºC, incubated with 20 μg/ml Alexa Fluor 633 succinimidyl ester at 37 ºC for 1 h, resuspended in PBS and 0.025% Tween 20 for use within 24 hr (Heung et al., 2015; Jhingran et al., 2016; Jhingran et al., 2012). To generate morphologically uniform heat-killed swollen conidia, 5Í10^6^/ml conidia were incubated at 37° C for 14 hours in RPMI-1640 and 0.5 μg/ml voriconazole and heat killed at 100 ºC for 30 minutes (Hohl et al., 2005). To infect mice with 30-60 million live or heat-killed *A. fumigatus* cells, conidia were resuspended in PBS, 0.025% Tween-20 at a concentration of 0.6-1.2Í10^9^ cells and 50 μl of cell suspension was administered via the intranasal route to mice anesthetized by isoflurane inhalation.

## EXPERIMENTAL DETAILS

### Analysis of *in vivo* and *in vitro* conidial uptake and killing

To analyze of conidia uptake and killing, FLARE conidia were used to infect mice (*in vivo*) or co-culture with BM leukocyte (*in vitro*). In data analyses for a given leukocyte subset, conidial uptake refers to the frequency of fungus-engaged leukocytes, i.e. the sum of dsRed+AF633+ and dsRed-AF633+ leukocytes. Conidial viability within a specific leukocyte subset refers to the frequency of leukocytes that contain live conidia (dsRed+AF633+) divided by the frequency of all fungus-engaged leukocytes (dsRed^+^AF633^+^ and dsRed^-^AF633^+^).

### *In vivo* Cell Depletion

To ablate specific cells, CCR2-DTR, CCR2-DTR, *Rag2*^-/-^*Il2rg*^-/-^, ROSA26-iDTR^Mrp8-Cre^, BDCA2-DTR, and non-transgenic littermate controls were injected i.p. with 10 ng/g body weight DT on Day −1, Day 0, and Day +2 pi (Espinosa et al., 2017; Swiecki et al., 2010), unless noted otherwise.

### Analysis of Infected Mice

Single cell lung suspensions were prepared for flow cytometry as described in (Hohl et al., 2009), with minor modifications. Briefly, perfused murine lungs were placed in a gentle MACS(tm) C tube and mechanically homogenized in 5 ml RPMI-1640, 10% FBS, and 0.1 mg/ml DNAse using a gentle MACS(tm) Octo Dissociator (Miltenyi Biotecc) in the absence of collagenase. Lung cell suspensions were lysed of RBCs, enumerated, and stained with fluorophore-conjugated antibodies prior to flow cytometric analysis on a BD LSR II or flow cytometric sorting on a BD Aria, flow plot analysis was performed using FlowJo v.9.6.6 software.

Neutrophils were identified as CD45+ CD11b+ Ly6Clo Ly6G+ cells, inflammatory monocytes as CD45+ CD11b+ CD11c-Ly6G-Ly6Chi cells, Mo-DCs as CD45+ CD11b+ CD11c+ Ly6G-Ly6Chi MHC class II+ cells, and pDCs as CD45+ CD11cint SiglecF-CD19-NK1.1-CD11b-B220+ SiglecH^+^ cells.

To analyze the lung fungal burden, perfused murine lungs were homogenized using a PowerGen 125 homogenizer (Fisher) in 2 mL PBS, 0.025% Tween-20, and plated on Sabouraud dextrose agar. To analyze cytokine levels by ELISA, whole lungs were weighed and mechanically homogenized in 2 mL PBS containing protease inhibitor. To analyze cytokine levels by qRT-PCR, total RNA from cells was extracted with TRIzol (Invitrogen). One to two micrograms of total RNA were reverse-transcribed using High-Capacity cDNA Reverse Transcription Kit (Applied Biosystems). TaqMan Fast Universal Master Mix (2×), TaqMan probes (Applied Biosystems) for each gene were used and normalized to glyceraldehyde-3-phosphate dehydrogenase. Gene expression was calculated using the *ΔΔ*Ct method relative to the naïve sample. For histology, perfused lungs were fixed in 4% paraformaldehyde, embedded in paraffin, sectioned in 4 μm slices, stained with hematoxylin and eosin (H&E) or modified Gomori methenamine silver (GMS), and digitally scanned using a Zeiss Mirax Desk Scanner. Images were captured from whole slide images, acquired with an Aperio ScanScope (Aperio Technologies) using 5× and 20× objectives at the Molecular Cytology Core Facility (MSKCC). BAL LDH and albumin levels were measured with a CytoTox96 non-radioactive cytotoxicity assay kit and a BCG albumin assay kit, respectively.

### Reactive Oxygen Production

Intracellular ROS levels were measured in cells using CM-H2DCFDA [5-(and 6-) chloromethyl-2,7-dichlorodihydrofluorescein diacetate, acetyl ester] as described in (Espinosa et al., 2017; Hackstein et al., 2012). Briefly, single cell lung suspensions were incubated with 1μM CM-H2DCFDA in Hanks’ balanced salt solution at 37º C for 45 min according to manufacturer’s instruction, and analyzed by flow cytometry.

## QUANTIFICATION AND STATISTICAL ANALYSIS

Data are representative of at least 2 independent experiments, as indicated. All results are expressed as mean (± SEM), unless stated otherwise. The Mann-Whitney test was used for comparisons of two groups, unless noted otherwise. The Kruskal-Wallis test was used for multi-group comparisons, unless noted otherwise. Survival data was analyzed by long-rank test. All statistical analyses were performed with GraphPad Prism software, v8.2.0.

## Graphical Abstract

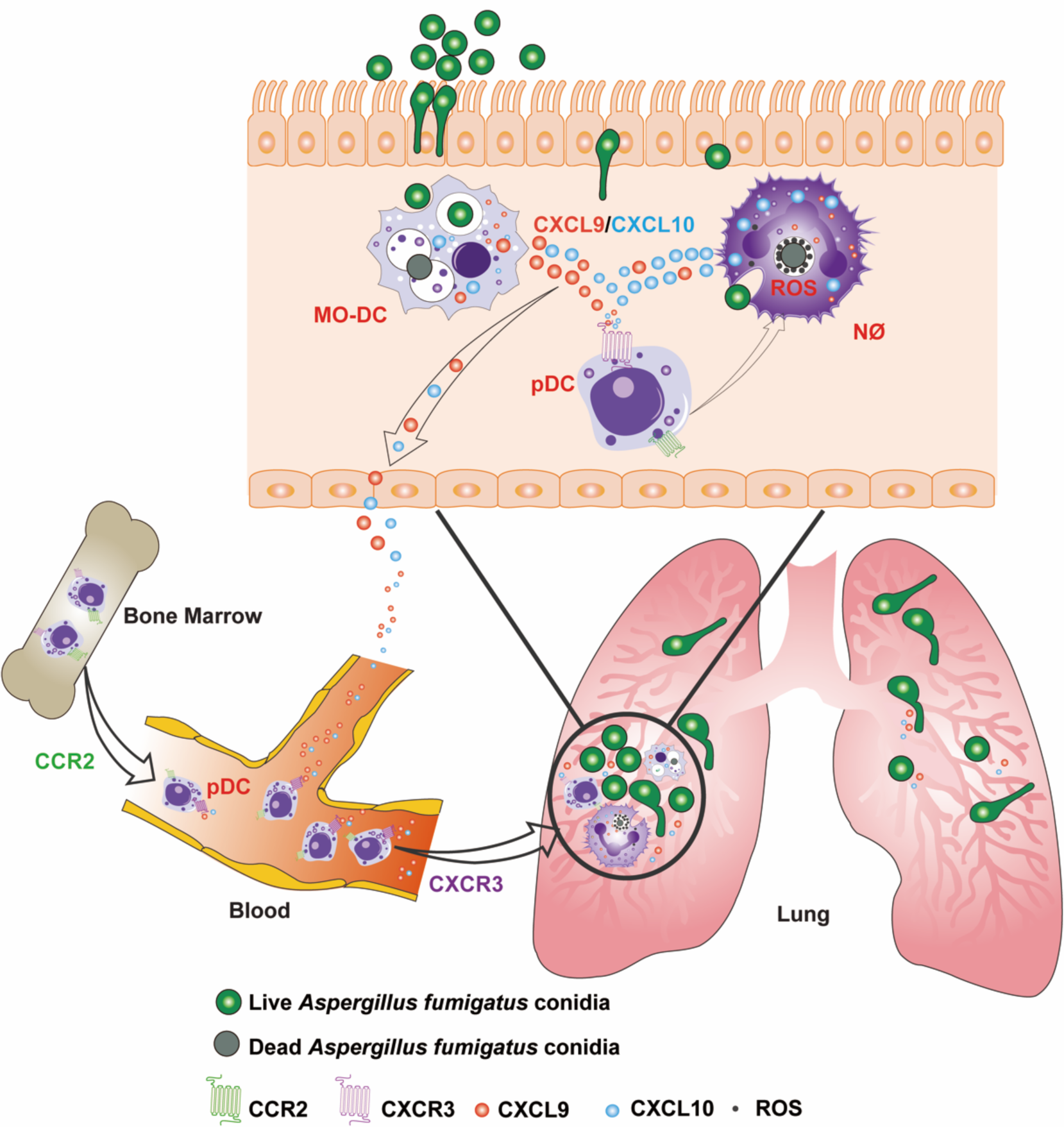

## SUPPLEMENTAL INFORMATION

**Figure S1.**
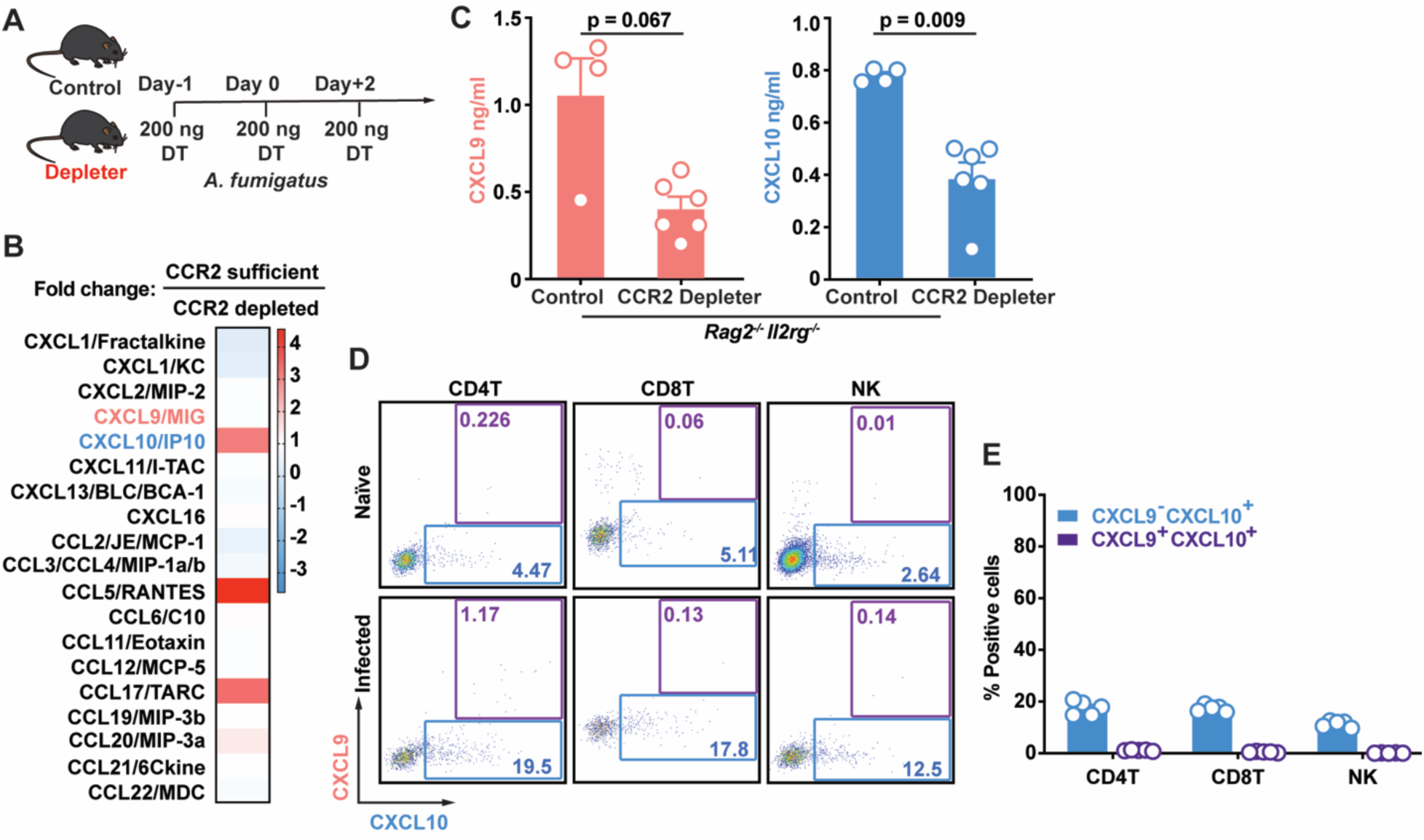
CCR2^+^ myeloid cells regulate CXCL9 and CXCL10 production during *A. fumigatus* infection. Related to Figure 1. (A) Experimental Scheme: Diphtheria toxin (DT) was administered intraperitoneally (i.p.) as indicated to ablate DTR^+^ cells in mouse strains that express the CCR2-DTR transgene (Depleter) or non-transgenic littermates that not do express the CCR2-DTR transgene (Control). (B) Fold change in lung chemokine levels in CCR2-sufficient (CCR2-DTR^-/-^ *Rag2*^*-/-*^*Il2rg*^*-/-*^) versus CCR2-depleted (CCR2-DTR^+/-^ *Rag2*^*-/-*^*Il2rg*^*-/-*^) mice that lack lymphoid lineage cells 36 h pi with 3 × 10^7^ CEA10 conidia, as measured by proteome profiler array (n = 2 per group, data pooled from 2 experiments). (C) Lung CXCL9 and CXCL10 levels in CCR2-DTR^+/-^ *Rag2*^*-/-*^*Il2rg*^*-/-*^ and littermates (CCR2-DTR^-/-^ *Rag2*^*-/-*^*Il2rg*^*-/-*^) (n = 4-6) at 48 h pi with 3 × 10^7^ CEA10 conidia. (D) Representative plots of RFP (CXCL9) and BFP (CXCL10) expression in indicated lung leukocytes isolated from Rex3 Tg → C57BL/6.SJL BM chimeric mice at baseline (naïve, top row) and 48 h pi with 3 × 10^7^ CEA10 conidia (infected, bottom row). The blue and purple gates indicate the frequency of BFP^+^ (CXCL9^+^) and BFP^+^RFP^+^ (CXCL9^+^ CXCL10^+^) cells, respectively. (E) The graphs indicate the frequency of CXCL9^-^ CXCL10^+^ and CXCL9^+^ CXCL10^+^ CD4^+^ T cells, CD8^+^ T cells and NK cells at 48 h pi. (C, E) Data are representative of two independent experiments. Dots represent individual mice and data are presented as mean ± SEM, (C) Statistical analysis: Mann-Whitney test.

**Figure S2.**
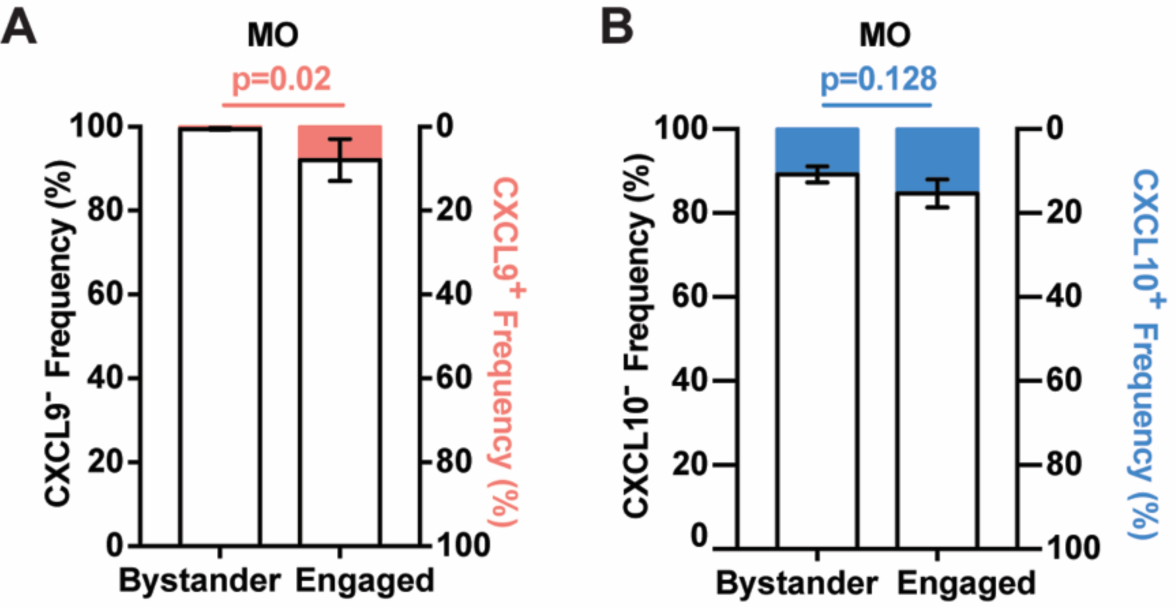
Monotye CXCL9 and CXCL10 expression during *A. fumigatus* infection. Related to Figure 2. (A) Proportion of RFP^+^ (CXCL9^+^; pink bar) and RFP^-^ (CXCL9^-^; white bar); and (B) BFP^+^ (CXCL10^+^; blue bar) and BFP^-^ (CXCL10^−^; white bar) expression in indicated bystander and fungus-engaged leukocytes isolated infected Rex3 Tg → C57BL/6.SJL BM chimeric mice (n = 7) with 3 × 10^7^ AF633-labeled CEA10 conidia. (A and B) Data are presented as mean ± SEM. Statistical analysis: Mann-Whitney test.

**Figure S3.**
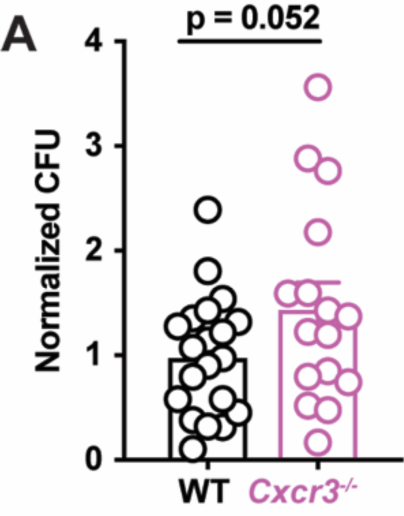
CXCR3 is critical for anti-*Aspergillus* defense. Related to Figure 3. (A) Normalized Lung CFUs in C57BL/6 (WT) and *Cxcr3*^-/-^ mice 72 h pi with 3 × 10^7^ CEA10 conidia. Dots represent individual mice and data were pooled from 2 independent experiments. Data are presented as mean ± SEM. Statistical analysis: Mann-Whitney test.

**Figure S4.**
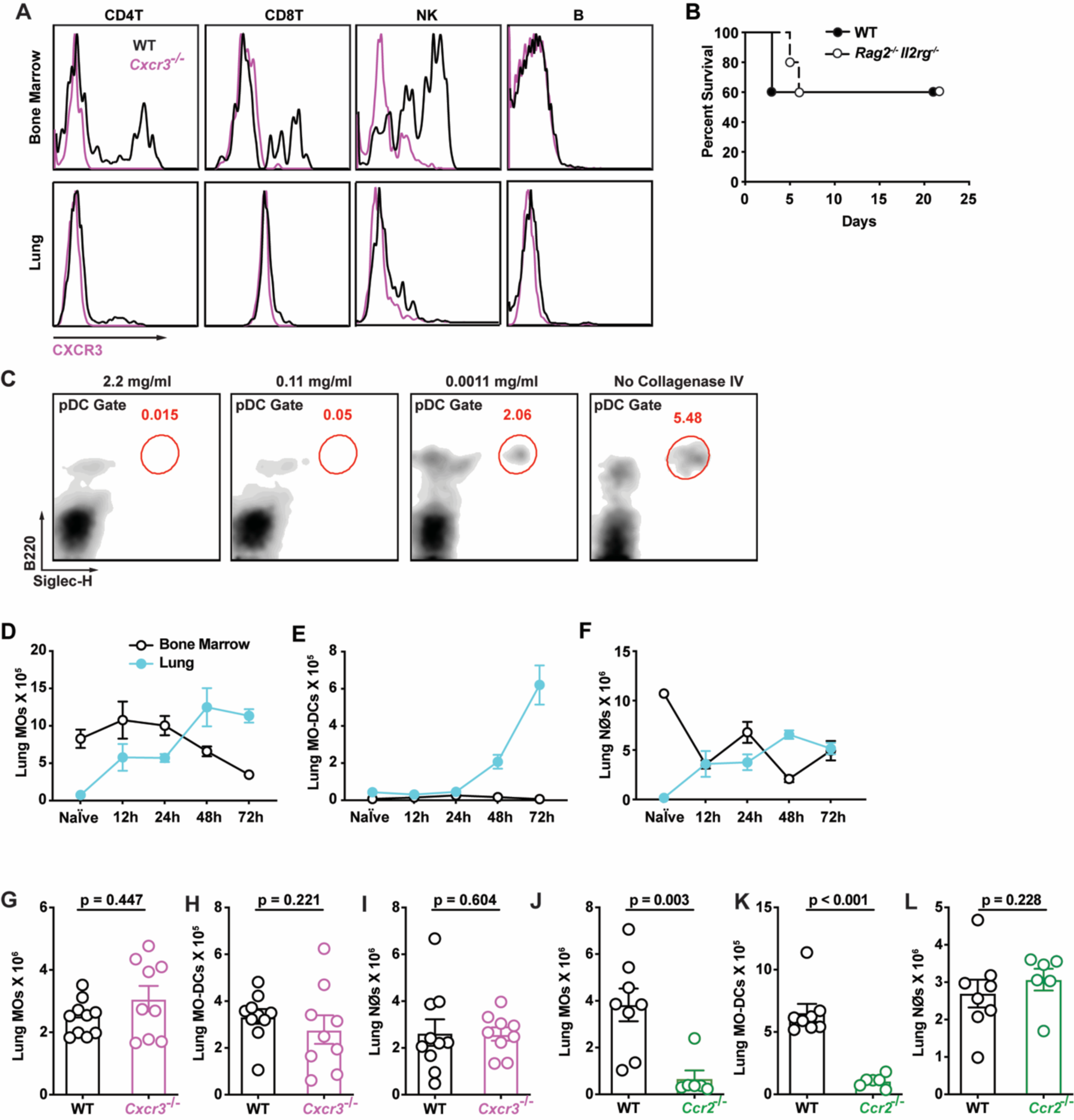
CXCR3 expression on lung leukocytes and pDC identification in lung digests. Related to Figure 4. (A) Representative CXCR3 surface expression in the indicated bone marrow (top row) and lung (bottom row) leukocytes that were isolated from C57BL/6 (WT, black lines; *Cxcr3*^+/+^) or *Cxcr3*^-/-^ mice (purple lines). (B) Kaplan-Meier survival of C57BL/6 (n = 5) and *Rag2*^-/-^*Il2rg*^-/-^ (n = 5) mice challenged with 4-8 × 10^7^ CEA10 conidia. (C) Representative flow cytometry plots of B220^+^Siglec-H^+^ lung pDCs with the indicated concentration of type IV collagenase included in lung preparations to obtain single cells for flow cytometric analysis. (D-F) Lung (blue filled dots) and bone marrow (open black dots) (D) monocyte, (E) Mo-DC, and (F) neutrophil numbers at baseline and indicated times pi with 3 × 10^7^ CEA10 conidia (n = 5). (G-H) Lung (G) monocyte, (H) Mo-DC, and (I) neutrophil numbers in C57BL/6 (WT; open black dots) and *Cxcr3*^-/-^ mice (open purple dots) at 72 h pi (n = 10). (J-L) Lung (J) monocyte, (K) Mo-DC, and (L) neutrophil numbers in C57BL/6 (WT; open black dots) and *Ccr2*^-/-^ mice (open green bars) at 72 h pi (n = 10). (D-L) Data are representative of two independent experiments. Dots represent individual mice and data are presented as mean ± SEM. Statistical analysis: Mann-Whitney test.

**Figure S5.**
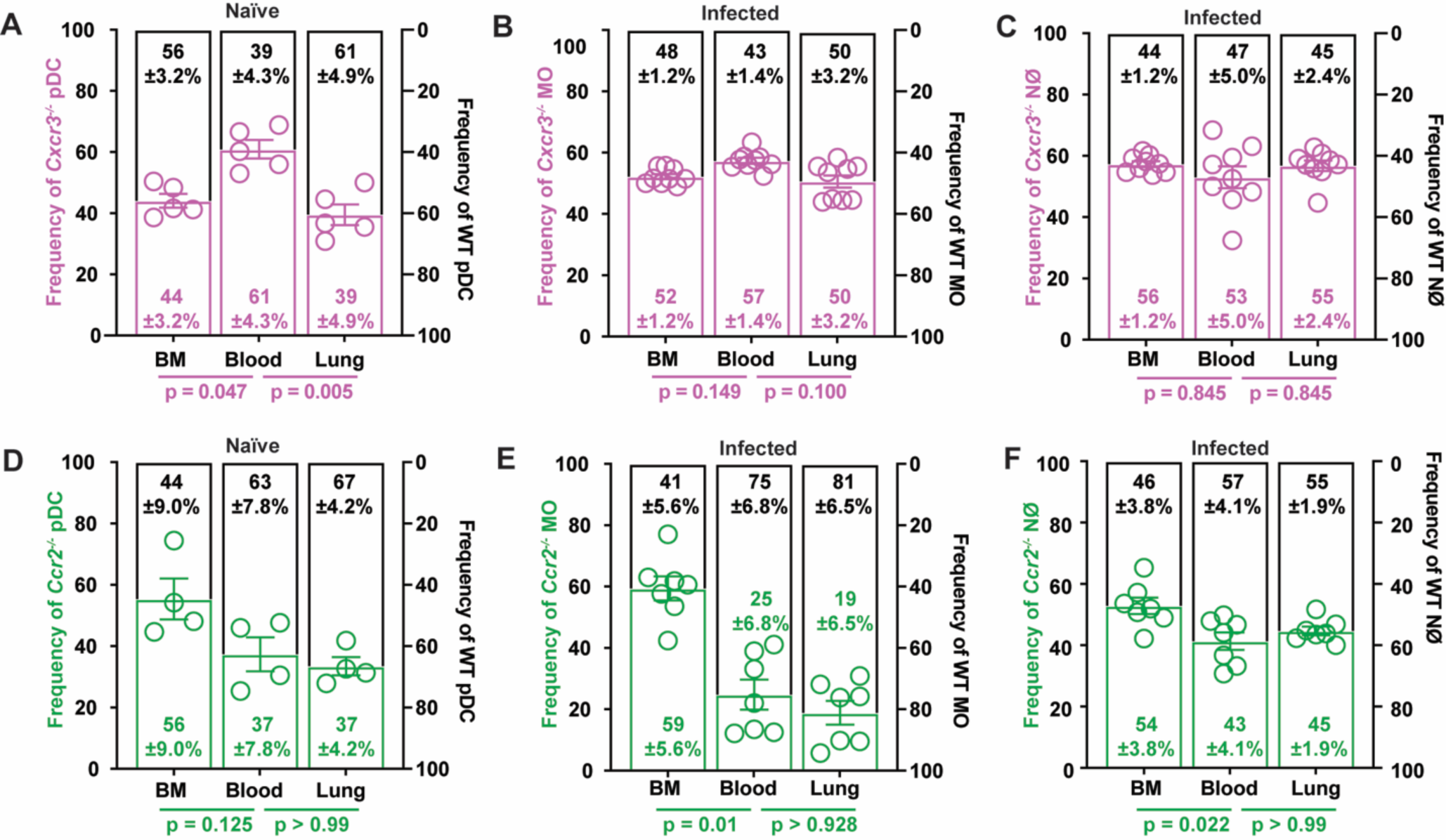
CXCR3 does not regulate the trafficking of lung monocytes, Mo-DCs, and neutrophils. Related to Figure 5. (A) Relative frequencies of *Cxcr3*^-/-^ (open purple bars) and *Cxcr3*^+/+^ (open black bars) pDCs in the BM, blood, and lung of mixed BM chimeric (1:1 mix of CD45.1^+^ *Cxcr3*^+/+^ and CD45.2^+^ *Cxcr3*^-/-^ BM cells → CD45.1^+^CD45.2^+^) mice at baseline. (B and C) Relative frequencies of *Cxcr3*^-/-^ (open purple bars) and *Cxcr3*^+/+^ (open black bars) (B) monocytes and (C) neutrophils in the BM, blood, and lung of mixed BM chimeric (1:1 mix of CD45.1^+^*Cxcr3*^+/+^ and CD45.2^+^*Cxcr3*^-/-^ BM cells → CD45.1^+^CD45.2^+^) mice 72 h pi.(D) Relative frequencies of *Ccr2*^-/-^ (open green bars) and *Ccr2*^+/+^ (open black bars) pDCs in the BM, blood, and lung of mixed BM chimeric (1:1 mix of CD45.1^+^*Cxcr3*^+/+^ and CD45.2^+^*Cxcr3*^-/-^ BM cells → CD45.1^+^CD45.2^+^) mice at baseline. (E-F) Relative frequencies of *Ccr2*^-/-^ (open green bars) and *Ccr2*^+/+^ (open black bars) (E) monocytes and (F) neutrophils in the BM, blood, and lung of mixed BM chimeric (1:1 mix of CD45.1^+^*Ccr2*^+/+^ and CD45.2^+^*Ccr2*^-/-^ BM cells → CD45.1^+^CD45.2^+^) mice 72 h pi. (A-F) Data were pooled from 2 or 3 independent experiments and presented as mean ± SEM, Statistical analysis: Mann-Whitney test.

**Figure S6.**
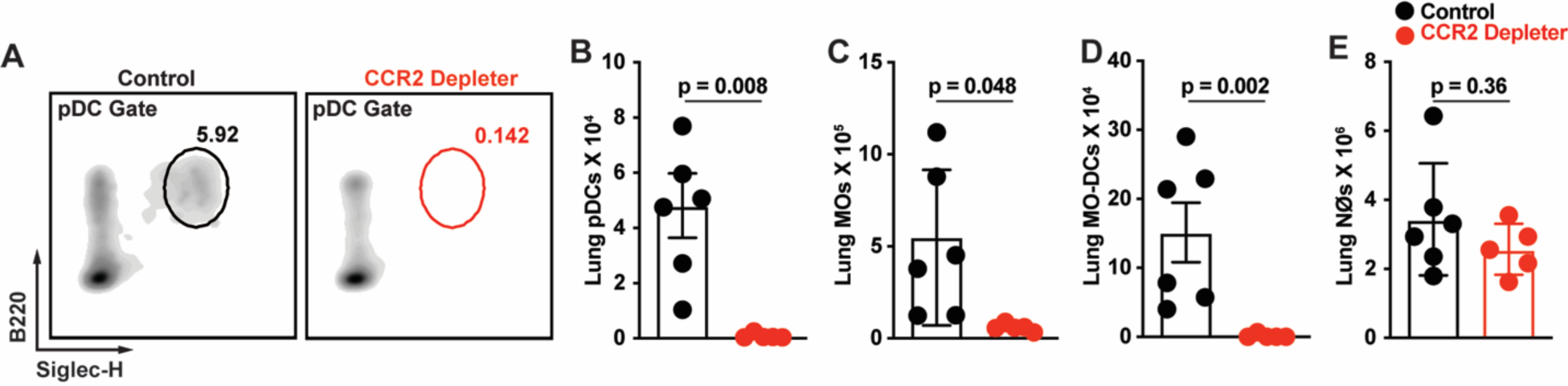
pDCs are depleted in CCR2 Depleter mice. Related to Figure 6. (A) Representative flow cytometry plots of lung B220^+^Siglec-H^+^ pDC, (B) lung pDC, (C) lung monocyte, (D) lung Mo-DC, and (E) lung neutrophil numbers in DT-treated CCR2 Depleter mice (CCR2-DTR^+/-^; red symbols) and non-Tg littermate controls (CCR2-DTR^-/-^; black symbols) at 72 h pi with 3 × 10^7^ CEA10 conidia. (B-E) Data were presented as mean ± SEM. Statistical analysis: Mann-Whitney test.

**Figure S7.**
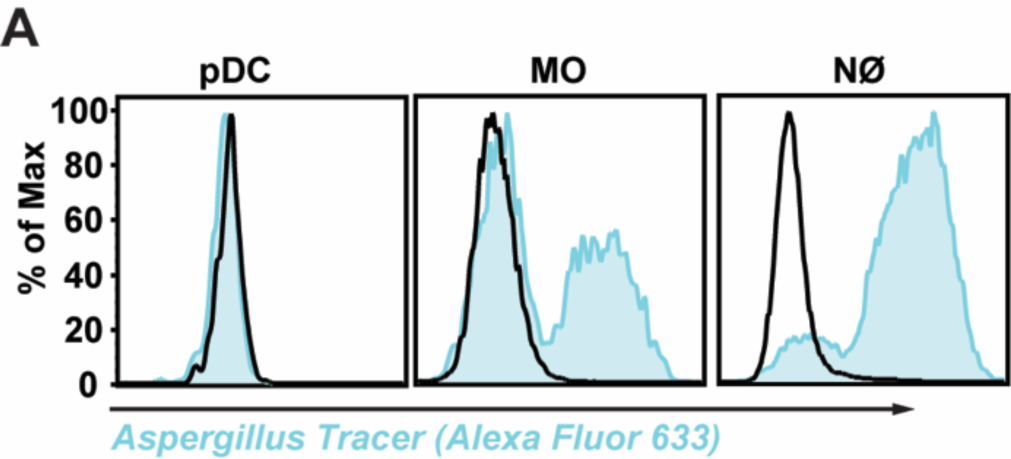
pDCs do not bind to or engulf *A. fumigatus* conidia. Related to Figure 7. (A) AF633 fluorescence intensity in indicated BM leukocytes co-cultured for 24 h with FLARE (blue line) or AF633-unlabeled conidia (MOI = 5).

