## Supplemental figure 1-7 for "During *Aspergillus* infection, neutrophil, monocyte-derived DC, and plasmacytoid DC enhance innate immune defense through CXCR3-dependent crosstalk"

SUPPLEMENTAL INFORMATION

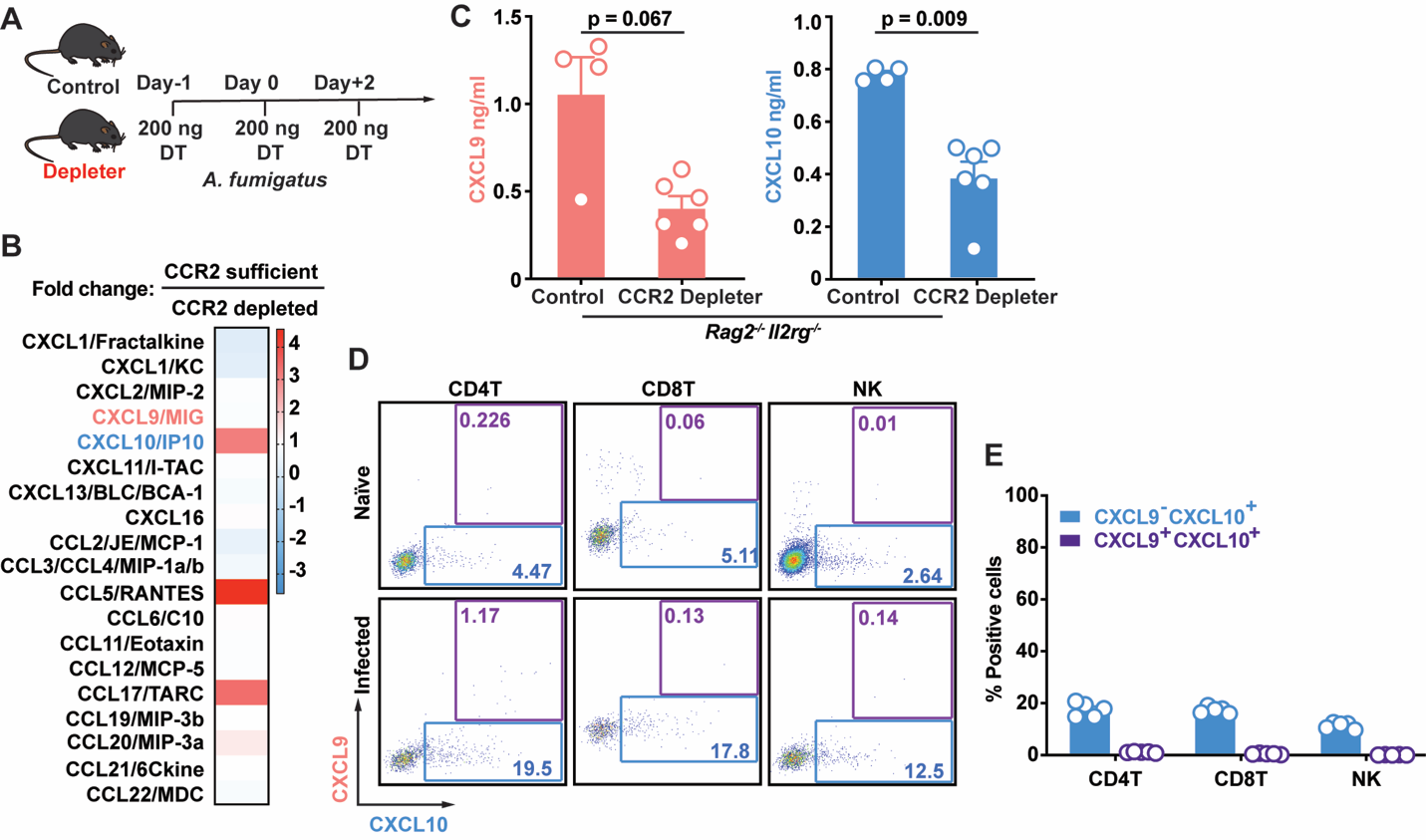

**Figure S1.** **CCR2^+^ myeloid cells regulate CXCL9 and CXCL10 production during *A. fumigatus* infection. Related to Figure 1.**

(A) Experimental Scheme: Diphtheria toxin (DT) was administered intraperitoneally (i.p.) as indicated to ablate DTR^+^ cells in mouse strains that express the CCR2-DTR transgene (Depleter) or non-transgenic littermates that not do express the CCR2-DTR transgene (Control).

(B) Fold change in lung chemokine levels in CCR2-sufficient (CCR2-DTR^-/-^ *Rag2^-/-^Il2rg^-/-^*) versus CCR2-depleted (CCR2-DTR^+/-^ *Rag2^-/-^Il2rg^-/-^*) mice that lack lymphoid lineage cells 36 h pi with 3 × 10^7^ CEA10 conidia, as measured by proteome profiler array (n = 2 per group, data pooled from 2 experiments).

(C) Lung CXCL9 and CXCL10 levels in CCR2-DTR^+/-^ *Rag2^-/-^Il2rg^-/-^* and littermates (CCR2-DTR^-/-^ *Rag2^-/-^Il2rg^-/-^*) (n = 4-6) at 48 h pi with 3 × 10^7^ CEA10 conidia.

(D) Representative plots of RFP (CXCL9) and BFP (CXCL10) expression in indicated lung leukocytes isolated from Rex3 Tg → C57BL/6.SJL BM chimeric mice at baseline (naïve, top row) and 48 h pi with 3 × 10^7^ CEA10 conidia (infected, bottom row). The blue and purple gates indicate the frequency of BFP^+^ (CXCL9^+^) and BFP^+^RFP^+^ (CXCL9^+^ CXCL10^+^) cells, respectively.

(E) The graphs indicate the frequency of CXCL9^-^ CXCL10^+^ and CXCL9^+^ CXCL10^+^ CD4^+^ T cells, CD8^+^ T cells and NK cells at 48 h pi.

(C, E) Data are representative of two independent experiments. Dots represent individual mice and data are presented as mean ± SEM, (C) Statistical analysis: Mann-Whitney test.

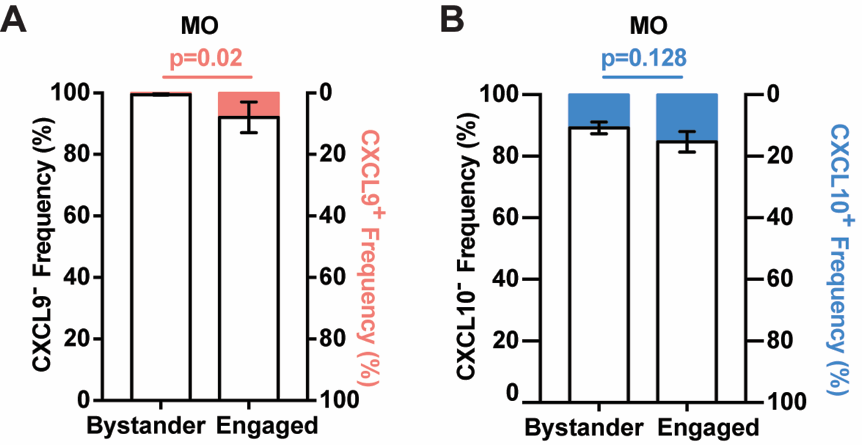

**Figure S2. Monotye CXCL9 and CXCL10 expression during *A. fumigatus* infection. Related to Figure 2.**

(A) Proportion of RFP^+^ (CXCL9^+^; pink bar) and RFP^-^ (CXCL9^-^; white bar); and (B) BFP^+^ (CXCL10^+^; blue bar) and BFP^-^ (CXCL10^-^; white bar) expression in indicated bystander and fungus-engaged leukocytes isolated infected Rex3 Tg → C57BL/6.SJL BM chimeric mice (n = 7) with 3 × 10^7^ AF633-labeled CEA10 conidia.

(A and B) Data are presented as mean ± SEM. Statistical analysis: Mann-Whitney test.

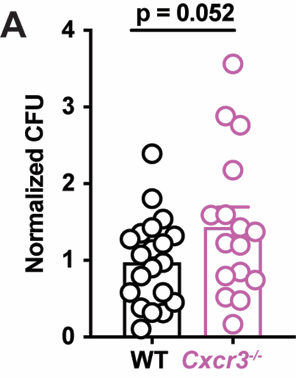

**Figure S3. CXCR3 is critical for anti-*Aspergillus* defense. Related to Figure 3.**

(A) Normalized Lung CFUs in C57BL/6 (WT) and *Cxcr3*^-/-^ mice 72 h pi with 3 × 10^7^ CEA10 conidia. Dots represent individual mice and data were pooled from 2 independent experiments. Data are presented as mean ± SEM. Statistical analysis: Mann-Whitney test.

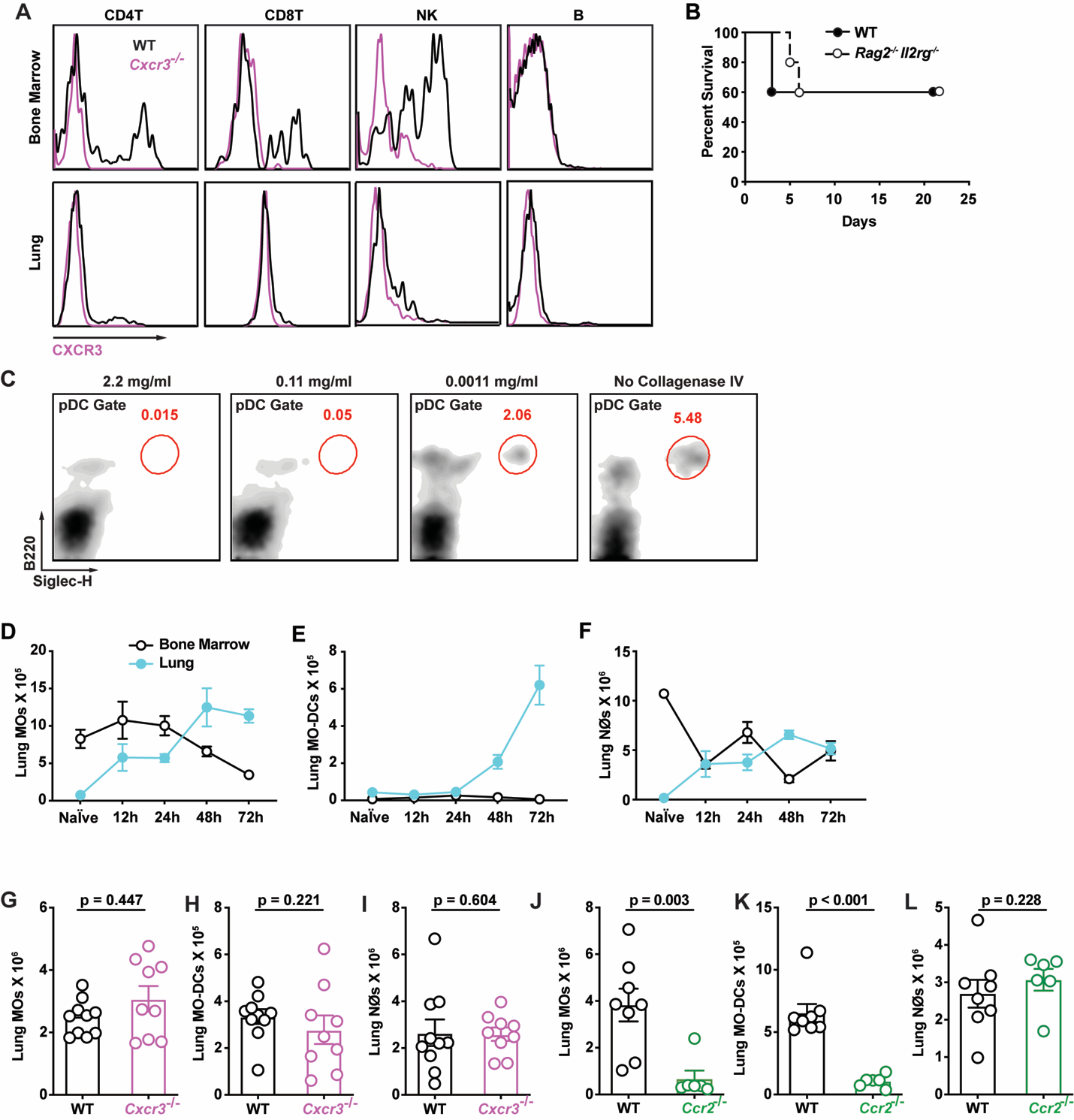

**Figure S4. CXCR3 expression on lung leukocytes and pDC identification in lung digests. Related to Figure 4.**

(A) Representative CXCR3 surface expression in the indicated bone marrow (top row) and lung (bottom row) leukocytes that were isolated from C57BL/6 (WT, black lines; *Cxcr3*^+/+^) or *Cxcr3*^-/-^ mice (purple lines).

(B) Kaplan-Meier survival of C57BL/6 (n = 5) and *Rag2*^-/-^*Il2rg*^-/-^ (n = 5) mice challenged with 4-8 × 10^7^ CEA10 conidia.

(C) Representative flow cytometry plots of B220^+^Siglec-H^+^ lung pDCs with the indicated concentration of type IV collagenase included in lung preparations to obtain single cells for flow cytometric analysis.

(D-F) Lung (blue filled dots) and bone marrow (open black dots) (D) monocyte, (E) Mo-DC, and (F) neutrophil numbers at baseline and indicated times pi with 3 × 10^7^ CEA10 conidia (n = 5).

(G-H) Lung (G) monocyte, (H) Mo-DC, and (I) neutrophil numbers in C57BL/6 (WT; open black dots) and *Cxcr3*^-/-^ mice (open purple dots) at 72 h pi (n = 10).

(J-L) Lung (J) monocyte, (K) Mo-DC, and (L) neutrophil numbers in C57BL/6 (WT; open black dots) and *Ccr2*^-/-^ mice (open green bars) at 72 h pi (n = 10).

(D-L) Data are representative of two independent experiments. Dots represent individual mice and data are presented as mean ± SEM. Statistical analysis: Mann-Whitney test.

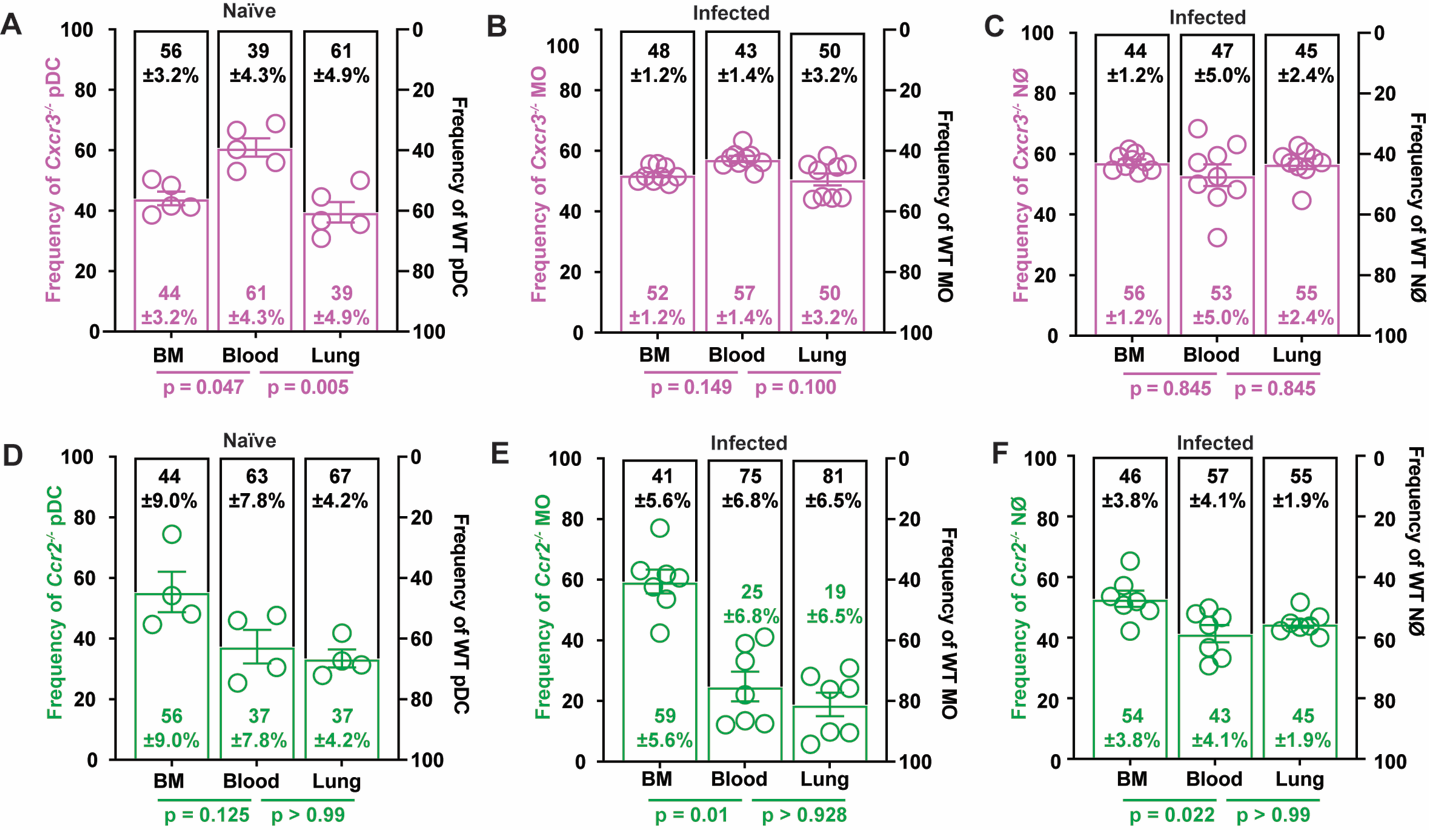

**Figure S5. CXCR3 does not regulate the trafficking of lung monocytes, Mo-DCs, and neutrophils. Related to Figure 5.**

(A) Relative frequencies of *Cxcr3*^-/-^ (open purple bars) and *Cxcr3*^+/+^ (open black bars) pDCs in the BM, blood, and lung of mixed BM chimeric (1:1 mix of CD45.1^+^ *Cxcr3*^+/+^ and CD45.2^+^ *Cxcr3*^-/-^ BM cells → CD45.1^+^CD45.2^+^) mice at baseline.

(B and C) Relative frequencies of *Cxcr3*^-/-^ (open purple bars) and *Cxcr3*^+/+^ (open black bars) (B) monocytes and (C) neutrophils in the BM, blood, and lung of mixed BM chimeric (1:1 mix of CD45.1^+^*Cxcr3*^+/+^ and CD45.2^+^*Cxcr3*^-/-^ BM cells → CD45.1^+^CD45.2^+^) mice 72 h pi.(D) Relative frequencies of *Ccr2*^-/-^ (open green bars) and *Ccr2*^+/+^ (open black bars) pDCs in the BM, blood, and lung of mixed BM chimeric (1:1 mix of CD45.1^+^*Cxcr3*^+/+^ and CD45.2^+^*Cxcr3*^-/-^ BM cells → CD45.1^+^CD45.2^+^) mice at baseline.

(E-F) Relative frequencies of *Ccr2*^-/-^ (open green bars) and *Ccr2*^+/+^ (open black bars) (E) monocytes and (F) neutrophils in the BM, blood, and lung of mixed BM chimeric (1:1 mix of CD45.1^+^*Ccr2*^+/+^ and CD45.2^+^*Ccr2*^-/-^ BM cells → CD45.1^+^CD45.2^+^) mice 72 h pi.

(A-F) Data were pooled from 2 or 3 independent experiments and presented as mean ± SEM, Statistical analysis: Mann-Whitney test.

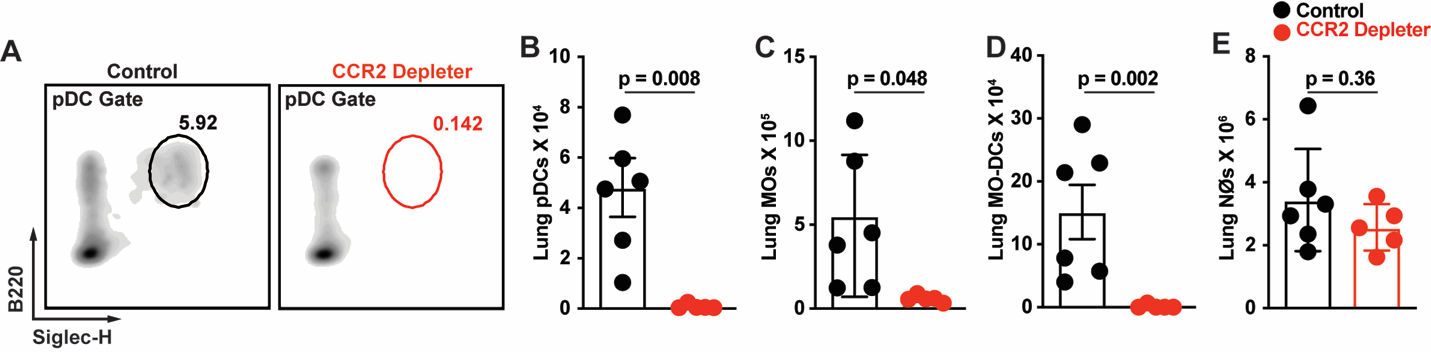

**Figure S6.** **pDCs are depleted in CCR2 Depleter mice. Related to Figure 6.**

(A) Representative flow cytometry plots of lung B220^+^Siglec-H^+^ pDC, (B) lung pDC, (C) lung monocyte, (D) lung Mo-DC, and (E) lung neutrophil numbers in DT-treated CCR2 Depleter mice (CCR2-DTR^+/-^; red symbols) and non-Tg littermate controls (CCR2-DTR^-/-^; black symbols) at 72 h pi with 3 × 10^7^ CEA10 conidia.

(B-E) Data were presented as mean ± SEM. Statistical analysis: Mann-Whitney test.

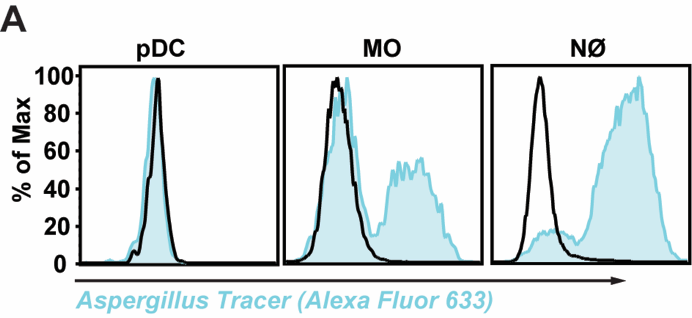

**Figure S7. pDCs do not bind to or engulf *A. fumigatus* conidia. Related to Figure 7.**

(A) AF633 fluorescence intensity in indicated BM leukocytes co-cultured for 24 h with FLARE (blue line) or AF633-unlabeled conidia (MOI = 5).
